# Brainwide blood volume reflects opposing neural populations

**DOI:** 10.1101/2025.07.01.662538

**Authors:** Agnès Landemard, Michael Krumin, Kenneth D Harris, Matteo Carandini

**Affiliations:** UCL Institute of Ophthalmology, University College London; UCL Queen Square Institute of Neurology, University College London

## Abstract

The supply of blood to brain tissue is thought to depend on the overall neural activity in that tissue, and this dependence is thought to differ across brain regions and across brain states. Studies supporting these views, however, measured neural activity as a bulk quantity, and related it to blood supply following disparate events in different regions. Here we measure fluctuations in neuronal activity and blood volume across the mouse brain, and find that their relationship is consistent across brain states and brain regions but differs in two opposing brainwide neural populations. Functional Ultrasound Imaging (fUSI) revealed that whisking, a marker of arousal, is associated with brainwide fluctuations in blood volume. Simultaneous fUSI and Neuropixels recordings showed that neurons that increase vs. decrease activity with whisking have distinct hemodynamic response functions. Their summed contributions predicted blood volume across states. Brainwide Neuropixels recordings revealed that these two opposing populations coexist in the entire brain. Their differing contributions to blood volume largely explain the apparent differences in blood volume fluctuations across regions. The mouse brain thus contains two neural populations with opposite relation to brain state and distinct relationships to blood supply, which together account for brainwide fluctuations in blood volume.

## Introduction

The blood supply to the brain is tightly correlated with neuronal activity^1–5^. This neurovascular coupling is usually described by a positive hemodynamic response function (HRF) that acts as a filter on the neural activity. Filtering the neural activity measured in a local region with the HRF provides a reasonable prediction of the blood supply in that local region^6–13^.

There is debate, however, about whether neuro-vascular coupling is consistent across brain regions^5,14^. Weaker HRFs have been reported in olfactory bulb^15^, somatosensory thalamus^16^, and frontal cortex^17^. Some studies even reported negative HRFs, such as in somatosensory cortex (ipsilateral to a sensory stimulus^18^) or in striatum^19^. Others argued that the HRFs are always positive (with decreases in blood supply stemming from decreases in firing rate^20^), and are consistent across brain regions^10,21^.

Moreover, neurovascular coupling might differ across brain states^5,22^. For instance, in mouse somatosensory cortex the correlation between blood supply and neural activity was reported to be stronger during sensory stimulation than at rest^23^, and stronger in NREM sleep than during wakefulness or REM sleep^24^. In macaques, moreover, neurovascular coupling was reported to be stronger when the animal rests^21^. Furthermore, changes in attention^25^ or anticipation^11,26^ seem to affect blood supply more than local neural activity. Perhaps these factors engage vascular mechanisms that do not depend only on neuronal firing^2^.

However, previous studies of neurovascular coupling summarized neural activity with bulk measures, which cannot distinguish the potentially different correlates of different neurons. Bulk measures of neural activity predict blood supply better than generic individual neurons^27^, but the pathways that couple neural activity to blood supply are complex and may well differ across cell types^5,28,29^. For example, this coupling might differ in neurons vs. glia^30^ or in excitatory vs. inhibitory^31–33^, and might be mediated by neuromodulators^13,34^ that are released by a minority of neurons. Moreover, combined electrophysiology and fMRI in cortex revealed that some neurons correlate positively with the local BOLD signal and others negatively^35^. Across the brain, there may be populations of neurons that have different patterns of activity and different coupling to blood supply, which would not be distinguishable in bulk measurements of neural activity.

Moreover, studies of neurovascular coupling measured neural activity in different regions in relation to different events. Studies focused on different regions, such as olfactory bulb^15^, somatosensory cortex^16^, frontal cortex^17^, or wider regions of the cortex^22^ measured neural activity with respect to different events, such as the delivery of an olfactory or somatosensory stimulus, or the onset of spontaneous running. Their results, therefore, are hard to compare and assemble into a unified view of neurovascular coupling across the brain.

To overcome these limitations, we measured brainwide blood volume and brainwide neural activity in relation to brainwide neural events. Functional Ultrasound Imaging^36^ (fUSI) revealed that whisking, a marker of arousal in rodents, is associated with stereotyped, brainwide changes in blood volume. Simultaneous fUSI and Neuropixels recordings^10^ revealed that neurons that increase vs. decrease activity with whisking^37–39^ have distinct hemodynamic response functions (HRFs). Their summed contributions predicted blood volume across states, from REM sleep to active wakefulness. Brainwide recordings of 18,791 neurons^40^ with Neuropixels probes^41^ revealed that these opposing populations coexist in every brain region. Their combined activity predicts the brainwide fluctuations in blood volume better than the bulk local activity. These results indicate that neurovascular coupling is consistent across brain regions and brain states, but is different in two brainwide neural populations with opposite relation to brain state.

## Results

To measure the changes in blood volume associated with spontaneous behaviours, we used functional Ultrasound Imaging^36^ (fUSI). In each session, we measured cerebral blood volume in a coronal imaging plane, simultaneously capturing its dynamics in many brain regions (Figure 1a). We varied the imaging plane across sessions, so that all planes together covered the posterior half of the cerebrum (cortex, hippocampal formation, striatum, and pallidum), all the interbrain (thalamus, hypothalamus, and zona incerta), most of the midbrain (including the substantia nigra and the superior colliculus), and part of the pons (Table S1). We aligned the imaging planes to the Allen Common Coordinate Framework Atlas^42^, assigned each voxel to a brain region, and computed the median activity across voxels within each region. Mice were head-fixed but free to run on a wheel. We monitored their behaviour with infrared cameras as they spontaneously went through varying states of arousal.

**Figure 1.**
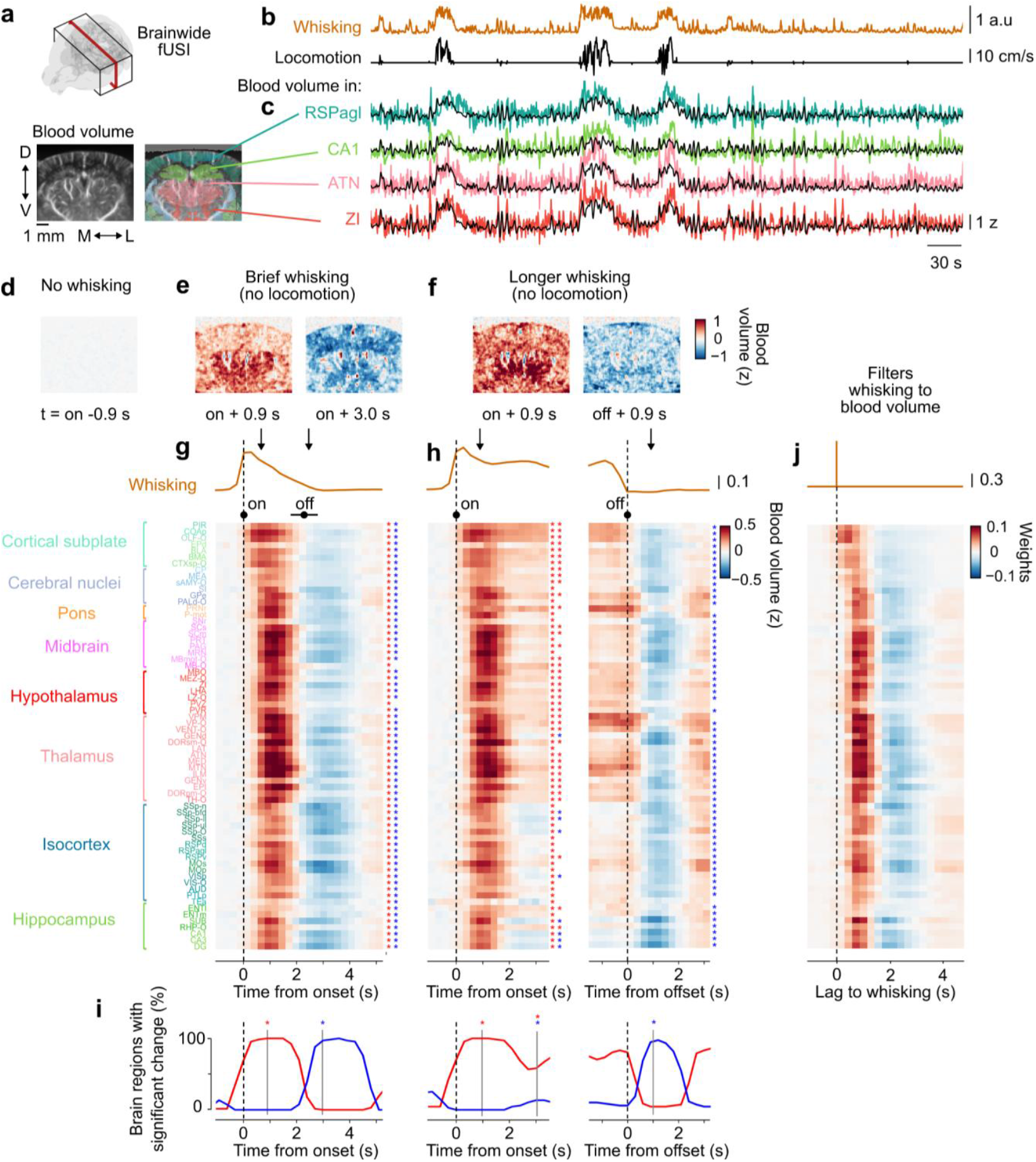
Brainwide blood volume is strongly modulated by arousal events. **a.** We measured blood volume with functional ultrasound imaging (fUSI), varying the coronal imaging planes (*red outline*) across sessions to image a volume of brain (*black outline*). Imaged volumes were aligned to the Allen Brain Atlas and voxels were averaged within brain regions (*colours*). **b**. Behavioural measurements — whisking and locomotion — from an example session. **c**. Corresponding fluctuations in blood volume in four example regions in isocortex (RSPagl), hippocampus (CA1), thalamus (ATN) and zona incerta (ZI, see Table S1 for abbreviations), showing predictions from whisking (*black*) with the filters in **j. d**. Average blood volume 0.9 s before the onset of whisking bouts in the example session, after subtracting the baseline (0.5–2 s before whisking onset). **e**. Same, for brief whisking bouts (1.3–3.5 s, without locomotion), 0.9 (*left*) and 3.0 s (*right*) after whisking onset. **f**. Same, for longer whisking bouts (> 3.5 s, without locomotion), 0.9 s after whisking onset (*left*), or offset (*right*). **g**. Average z-scored changes in blood volume across all sessions and mice, in each imaged region during brief whisking bouts (1.3–3.5 s) without locomotion. *Arrows* indicate the times of the images in **e**. *Black dots* below whisking indicate bout onset and average offset ± 1 s.d. Results are averaged first within mice and then across mice (number of sessions and events indicated in Figure S1). *Asterisks* denote significant increases (*red*) and decreases (*blue*) at 0.9 s and 3.0 s (permutation test). **h**. Same, for longer whisking bouts without locomotion (> 3.5 s, average = 12.1 s), aligned to bout onset (*left*) and offset (*right)*. See Figure S4 for whisking bouts associated with locomotion. **i**. Fraction of regions with significant change in blood volume at each timepoint, determined using a session permutation test. Dotted lines indicate the timepoints used to measure significance in g–h. **j**. Best-fitting filters to predict blood volume from whisking, for all brain regions (Figure S2, Figure S3). These filters model changes in blood volume for a notional delta function (impulse) of whisking (*top*). Example predictions are shown in **a** (*black*).

Increases in arousal marked by bouts of whisking^38,43,44^ were associated with widespread and dynamic changes in blood volume. Whisking, often accompanied by locomotion^45^(Figure 1b), correlated with fluctuations in blood volume in multiple brain regions (Figure 1c). In a typical session, brief bouts of whisking led to a widespread increase in blood volume followed by a decrease (Figure 1d, e). Longer bouts led to a similar pattern, with decreases in blood volume occurring at whisking offset (Figure 1f).

Brainwide measurements of blood volume established that correlates of whisking were present in the whole brain. We repeated our fUSI measurements in 5 mice and 138 sessions (101 hours of imaging, Figure S1) across multiple coronal planes, and we combined the results for each brain region. Brief whisking bouts (1.3–3.5 s, without locomotion), led to an almost simultaneous increase in blood volume in the entire brain, followed by an almost simultaneous decrease around 2 s after the bout began (Figure 1g). Longer whisking bouts led to a longer brainwide increase in blood volume, followed by a similar simultaneous decrease when the bout stopped (Figure 1h). These effects were not explainable by artefactual effects of brain movement (Methods). They were significant in practically all brain regions (Figure 1g–i), with occasional regional differences in amplitude and dynamics. For example, hippocampus and some regions of isocortex only showed a transient increase lasting ~2 s, followed by a decrease even while whisking continued; ventral regions such as amygdala and olfactory areas tended to activate earlier than the others at the onset of whisking bouts (Figure 1g, h).

Whisking was a strong predictor of fluctuations in blood volume across arousal states and brain regions. We used linear regression to obtain filters (Figure 1j) that predict each region’s blood volume from whisking (Figure S2). The obtained filters were biphasic and highly similar across regions (Figure 1j). Their predictions accounted for a large fraction of the variance in blood volume across states of arousal, from stillness to locomotion, and across multiple regions (Figure 1c, Figure S3). Thus, brainwide fluctuations in blood volume can be predicted by applying a biphasic filter to an arousal factor indexed by whisking, and the predictions based on whisking encompass the correlates of running (Figure S4).

We next investigated the neural basis of the whisking-related changes blood volume. We used simultaneous recordings of hundreds of neurons in visual cortex and hippocampus with Neuropixels and blood volume with fUSI^10^ in head-fixed mice that sat on a platform (Figure 2a). We binned spikes to the same temporal resolution as the fUSI data (300 ms time bins) and computed the overall firing rate of the neurons, along with whisking and blood volume in visual cortex (Figure 2b).

**Figure 2.**
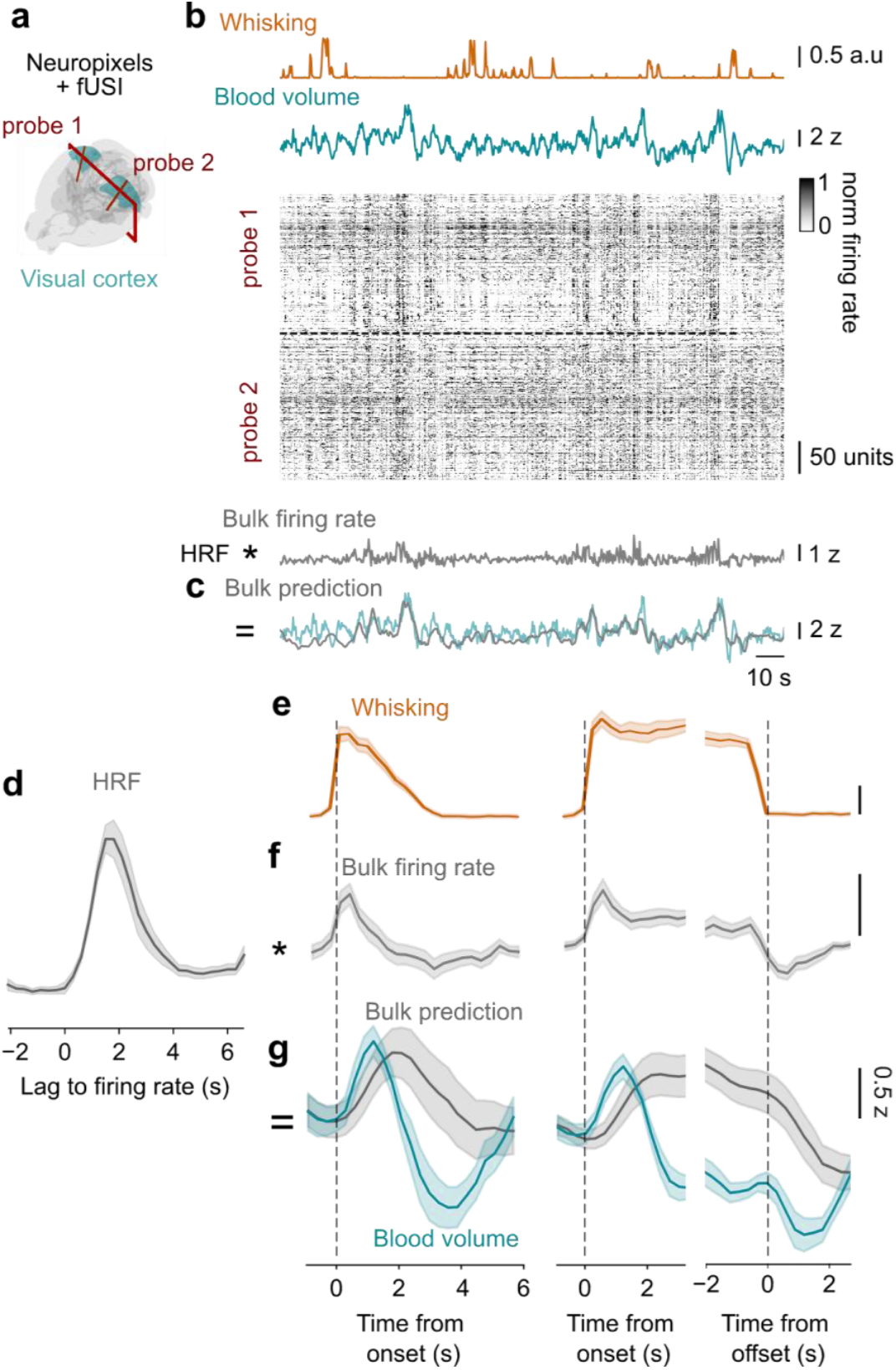
Bulk firing rate fails to predict changes in blood volume with whisking events. **a.** While imaging blood volume with fUSI, we used Neuropixels probes to record from hundreds of neurons in visual cortex and hippocampus^10^. **b**. Snippet from a typical recording session in visual cortex, with whisking (*gold*), blood volume (*cyan*) along with the normalized firing rate of all neurons recorded bilaterally across two probes in visual cortex. The firing rate for each neuron is divided by the 95^th^ percentile of firing rate across time. Neurons are sorted by depth on each probe. Bulk firing rate (*grey*) is obtained by taking the average over all neurons. **c**. The filtered bulk firing rate (*grey*) to predict blood volume (*cyan*) throughout the session. **d**. The best-fitting filter to predict blood volume from bulk firing rate. **e**. We identified whisking bouts and looked at brief (*left*) and longer whisking bouts, aligned to onset (*middle*) or offset (*right*). **f**. Average changes in bulk firing rate during whisking events. **g**. Average changes in blood volume during whisking events (*cyan*), and prediction from bulk firing rate in visual cortex (*grey*). **d**–**g** show results averaged across all recording sessions in visual cortex (see Figure S8 for similar results in hippocampus). Shaded area shows the s.e. across sessions.

We first attempted to predict the changes in blood volume from the local bulk firing rate, and failed to capture the correlates of whisking. Following established practice^6–12,46^, for each recording session, we fit a linear filter to predict blood volume from the average firing rate. As expected^10^, the best-fitting filter provided good overall predictions of blood volume (Figure 2c). Across sessions the best-fitting filter was monophasic and peaked 1.8 s after neural activity (Figure 2d), consistent with the standard HRFs that have been obtained from bulk neural activity in mice^10,47,48^. However, closer inspection revealed that the prediction tended to be sluggish, failing to capture the fastest dynamics of spontaneous blood volume (Figure 2c). This limitation was particularly clear during whisking events. The bulk firing rate measured during whisking events showed an increase followed by a smaller decrease (Figure 2e, f). This slightly biphasic response is in principle consistent with the biphasic response in the simultaneously measured blood volume. However, convolution with the HRF (Figure 2d) smoothed this feature out, failing to reproduce the biphasic changes in blood volume seen during whisking events (Figure 2g). Thus, a linear filter on average neural activity could not properly account for changes in blood volume during whisking events. To account for brainwide blood volume, it might be necessary to consider more than one dimension of neural activity.

Closer inspection of the neural data revealed two populations of neurons with opposite modulation by arousal. Consistent with previous measurements^38^, we found that neurons differed substantially in their relation to arousal: some neurons correlated positively with whisking and others correlated negatively (Figure 3a-c). We refer to these two opposing populations as Arousal+ and Arousal– neurons. These populations likely correspond to the “positive-delay” and “negative-delay” neurons that have been described in multiple brain regions in relation to global neural events^49^ associated with pupil dilations and running^50^ (Figure 3c, Figure S5). In the cerebral cortex, the Arousal+ neurons were less likely to be putative inhibitory (3.4%) than Arousal– neurons (5.8%, p = 0.03, t-test across sessions), and more likely to reside in the infragranular than the supragranular layers (44 vs. 23%, p = 0.027, t-test across sessions).

**Figure 3.**
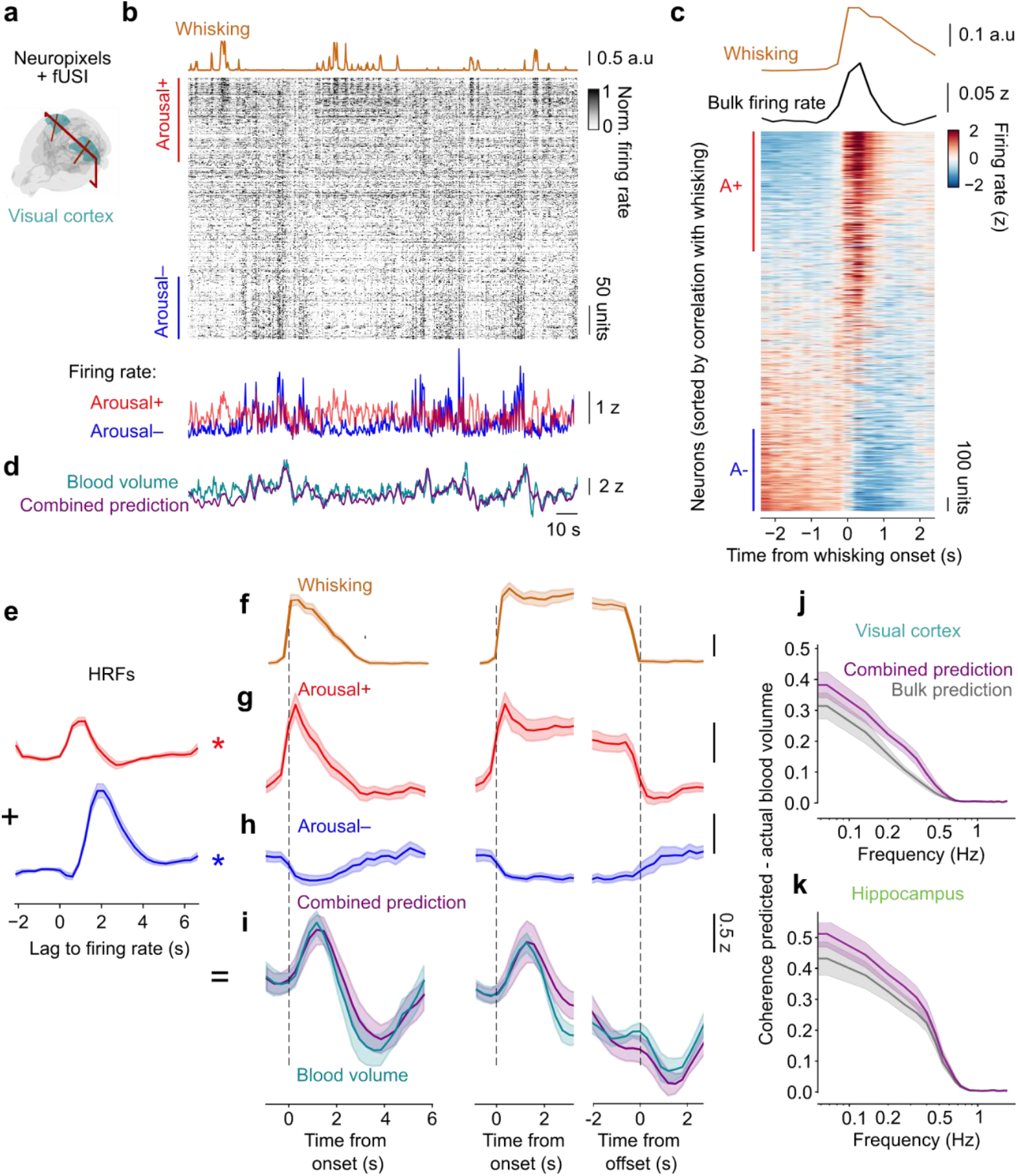
The firing rate of two arousal–defined neural populations best predicts blood volume. **a.** We analysed the same simultaneous Neuropixels-fUSI recordings as in Figure 2. **b**. *Top:* Same snippet of recording as in Figure 2, with neurons sorted by correlation with whisking. The firing rate for each neuron is divided by the 95^th^ percentile of firing rate across time. Bars indicate Arousal+ neurons (*red*, correlation with whisking > 0.05), and Arousal– neurons (*blue*, correlation with whisking < −0.05). *Bottom:* Average firing rate for Arousal+ neurons (*red*), and Arousal– neurons (*blue*). The firing rate of each neuron is z-scored before averaging. **c**. Firing rate of all neurons recorded in visual cortex across all sessions, aligned to and averaged across whisking events. The traces (*top*) show the corresponding average whisking (*gold*), and bulk firing rate (*black*). Each row is z-scored. Neurons are sorted by their correlation with whisking. **d**. Same experiment as **b**, showing the simultaneously measured blood volume (*cyan*), and the combined prediction from the firing of Arousal+ and Arousal– neurons (*purple*). **e**. The best-fitting filters to predict blood volume from Arousal+ (*red*) and Arousal– (*blue*) neurons. **f**. Traces of whisking for brief (*left*) and longer whisking events, aligned to onset (*middle*) or offset (*right*). See Figure S7 for the briefest whisking bouts. **g**. Average changes in firing rate for Arousal+ neurons during whisking events. **h**. Same, for Arousal– neurons. **i**. Average change in blood volume during whisking events (*cyan*), and prediction from Arousal+ and Arousal– neurons in visual cortex (*purple*). Data in **e**–**i** is averaged across all recording sessions in visual cortex (see Figure S8 for results in hippocampus). **j**. Coherence between blood volume and the predictions from bulk firing rate (*grey*, as in Figure 2) or from Arousal+ and Arousal– neurons (*purple*), in visual cortex. **k**. Same, for recordings in hippocampus. In **e-k**, shaded area shows the s.e. across sessions.

We thus asked whether considering these opposing neural populations might yield better predictions of blood volume. We designed a model where the blood volume in a region reflects not just the bulk neural activity but rather the activity of the two opponent populations of Arousal+ and Arousal– neurons, each acting with its own HRF. For each session, we filtered the activity of Arousal+ and Arousal– neurons with the best-fitting filters to obtain a prediction of blood volume (Figure 3d). The resulting filters were positive and monophasic, but with different peak times and widths (Figure 3e, Figure S6): the filter for Arousal+ neurons peaked earlier (1.2 s after neural activity) and was narrower (1.2 s full width at half height), whereas the filter for Arousal– neurons peaked later (2.1 s after neural activity) and was wider (1.8 s). They also differed in size (integral over time), with the filter for Arousal– neurons 1.1–2.4 times larger than for Arousal+ neurons (95% confidence interval, mean = 1.8, n = 10 sessions, p = 2e-4).

This combined model provided better estimates of the local blood volume than the traditional bulk model. Compared to the bulk prediction (shown in Figure 2c), the combined prediction was better able to track the observed fluctuations in blood volume, even in the absence of whisking (Figure 3d). Moreover, it provided a good account of the blood volume fluctuations associated with whisking events. At whisking onset, the activity of Arousal+ neurons increased (Figure 3f, g), while the activity of Arousal– neurons decreased (Figure 3h). The combined prediction from these two populations captured the characteristic biphasic changes in blood volume seen during whisking events (Figure 3i, Figure S7). The prediction first increased due to the increase in activity of Arousal+ neurons and later decreased due to the delayed decrease in activity of Arousal– neurons. More generally, regardless of whisking events, the firing rates of the two opposing populations provided better estimates of blood volume than the bulk firing rate (Figure 3d): their predictions had higher coherence with the measured blood volume at all frequencies, both in visual cortex (p = 0.003 across frequencies, Figure 3j) and in hippocampus (p = 0.0004, Figure 3k, Figure S8). This improvement was present during both spontaneous activity and visual stimulation (Figure S9).

The changes in blood volume seen in visual cortex and hippocampus during whisking events, therefore, are best explained by the activity of two opposing populations of neurons, each with its own hemodynamic response function. The improved performance of the predictions from the two opposing populations cannot be explained by having more free parameters because the fits were performed with cross-validation (see Methods). Moreover, the improvement was specific to those two populations of neurons: it was not present when using either one alone, or when using two random populations of neurons (Figure S10). Finally, we assessed whether a biphasic HRF would provide a good account of blood volume based on average firing rate only. To do so, we computed a biphasic HRF from the difference between the HRF for Arousal+ neurons (Figure 3e, top), and the HRF for Arousal– neurons (Figure 3e, bottom), with different weights. Applying these filters to the average firing rate was not sufficient to recapitulate the changes in blood volume during whisking events nor to reach similar levels of accuracy as the model from the two populations (Figure S11). Thus, the two populations of Arousal+ and Arousal– neurons provide a superior account of blood volume, which cannot be achieved from bulk firing rate alone.

The only way in which we could make the predictions of bulk firing rate approach the quality of the combined model was by pairing it with an additional predictor: the pupil dilation. Pupil dilation is a strong marker of arousal^12,51–54^ and an excellent predictor of blood volume in the visual cortex^47^. We tested a model that used both bulk firing rate and pupil dilation as predictors (Figure S10). In visual cortex, this model was superior to the prediction from bulk activity alone (p = 7e-3) and similar to our combined model based on Arousal+ and Arousal– populations (p = 0.35). In hippocampus however, it showed no significant improvement over the bulk model (p = 0.09) and was inferior to the combined model (p = 3e-3). Its performance was relatively worse at frequencies around 0.2–0.7 Hz, presumably because it lacked the fast dynamics of Arousal+ and Arousal– populations around whisking events.

The combined model provided good estimates of blood volume across the full range of brain states, from REM sleep to active wakefulness. During a typical session, the mouse spontaneously drifted across brain states (Figure 4a, b), marked by characteristic^12,55^ variations in pupil dilation, whisking, eye movements, and eye position (Figure 4c). The pupil is most constricted in REM and dilates with arousal^12,56^. Eye movements^57,58^ are characteristically frequent in REM, minimal in NREM and quiet wakefulness, and pronounced in active wakefulness. Similarly, whisking has small amplitude in REM^39^, is rare in NREM, and maximal in active wakefulness. Eye position, finally, is downcast and nasal during REM^12,58^. Across this range of brain states, the populations of Arousal+ and Arousal– neurons tended to show opposing activity patterns, which in combination provided good estimates of the local blood volume (Figure 4b). Consistent with their names, Arousal-neurons fired most during sleep and least during wakefulness, and Arousal+ neurons did the opposite (Figure 4d). Because the model weighs Arousal– neurons ~1.8 times more than Arousal+ neurons, it correctly predicts that blood volume is maximal during REM^24,59^ (when Arousal– neurons fire the most), low during quiet wakefulness, and intermediate during active wakefulness (when Arousal+ neurons fire the most) (Figure 4e). By contrast, the bulk model performed well during sleep^24^ but incorrectly predicted that blood volume should be similar in REM and active wakefulness (Figure 4e). Because the model assigns slightly delayed hemodynamic response function to Arousal– neurons, it also accounts for the longer delay^24^ in the correlation between (bulk) neural activity and blood volume seen in sleep relative to alertness, which could not be predicted by the bulk model (Figure 4f). Overall, the combined model performed as well as the bulk model in sleep states and was significantly superior to it in the awake states (Figure 4g). The inferior performance of the bulk model could not be rescued by allowing it to take different HRFs in different states (Figure 4g): it lies in its failure to distinguish the contributions of the two classes of neurons.

**Figure 4.**
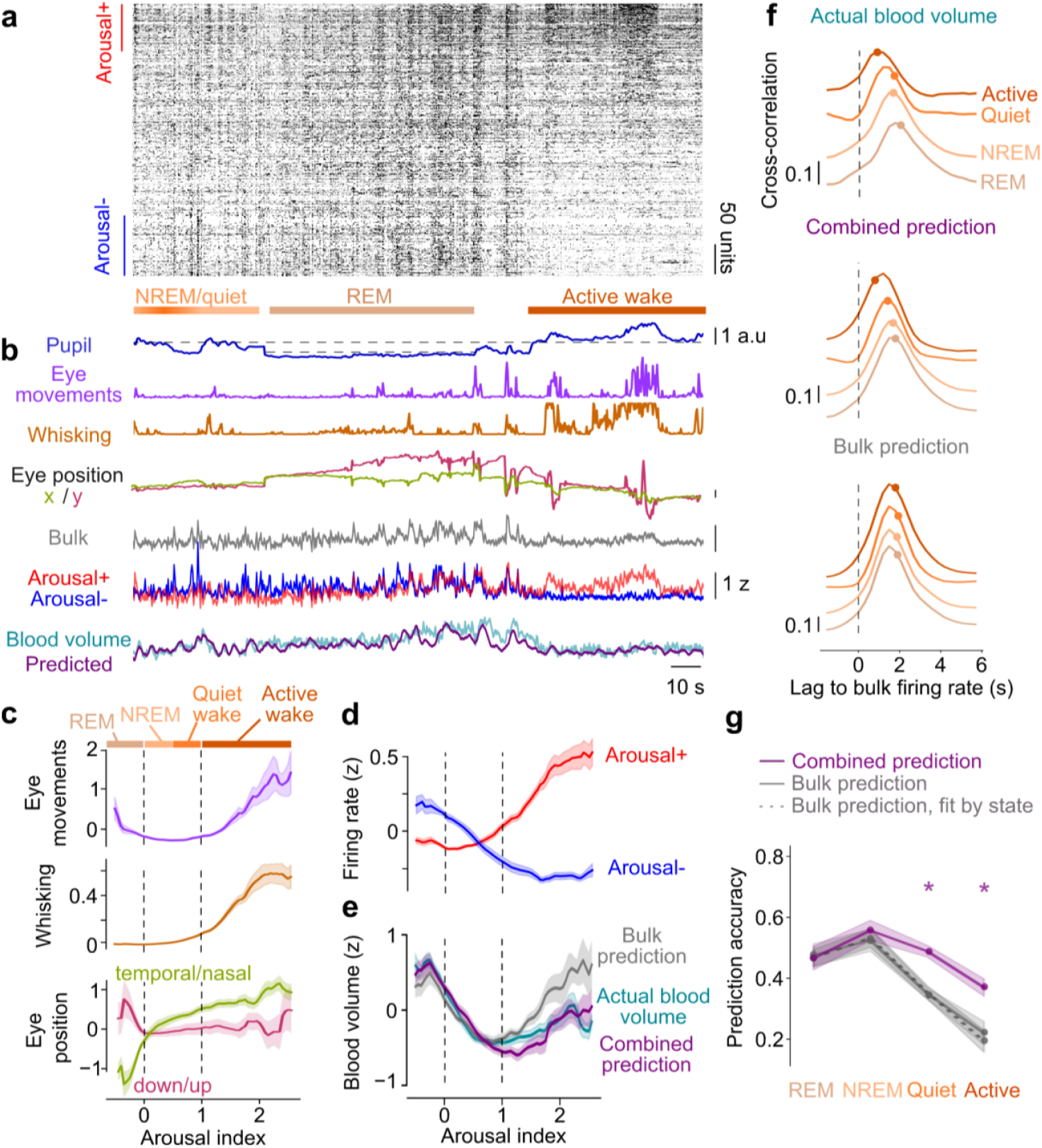
The firing rate of the two populations, but not the bulk firing rate, accounts for blood volume across states. **a.** Snippet from a simultaneous Neuropixels-fUSI recording session in visual cortex, as in Figure 3b. **a**. Normalized firing rate of all neurons recorded in visual cortex sorted by correlation with whisking. The firing rate for each neuron is divided by the 95^th^ percentile of its firing rate across time. Bars indicate Arousal+ neurons (*red*, correlation with whisking > 0.05), and Arousal– neurons (*blue*, correlation with whisking < −0.05). **b**. We use a normalized version of pupil size for each recording session to define an arousal index, from which we derive four states: putative rapid eye-movement sleep (REM), putative non-rapid-eye-movement sleep (NREM), quiet wake, and active wake. This snippet shows an episode of putative REM, with very constricted pupil, presence of eye movements, downcast and nasal eye position, increased activity of both Arousal+ and Arousal– neurons, as well as elevated blood volume. **c**. Behavioural metrics as a function of arousal index, defined as a scaled version of pupil size in 3 s intervals. 0 corresponds to the transition to putative REM, with increased eye movements, and characteristic eye positions. 1 corresponds to the transition to active wake, with increased whisking and eye movements. NREM and quiet wake are defined as transition states between these extremes. **d**. Firing rate of Arousal+ (*red*) and Arousal– (*blue*) neurons as a function of arousal index. **e**. Actual blood volume (*cyan*) and predictions from bulk (*grey*) and combined (*purple*) models, as a function of arousal index. **f**. Cross-correlation between bulk firing rate and actual blood volume, blood volume predicted from the bulk or combined model, for each state. Dots indicate the centre of mass of the cross-correlations for each state. **g**. Prediction accuracy, computed as the Pearson correlation between actual and predicted blood volume, depending on state. State is computed based on arousal index in 10 s intervals. *Asterisks* indicate significant differences between combined and bulk prediction (paired t-test, p < 0.05, n = 10 recording sessions).

To test whether a similar model applied in other brain regions, we turned to recordings with Neuropixels probes^41^ across the entire mouse brain^40^. In these recordings, each brain region is covered by multiple probe insertions. We focused on the periods after the end of decision-making task sessions, when the mice rested passively (n = 324 recordings from 108 mice), and exhibited fluctuations in arousal marked by whisking. Again, we binned spikes to the same temporal resolution as the fUSI data (300 ms time bins), and we assigned neurons to brain regions using the same mapping (e.g., Figure 5a).

**Figure 5.**
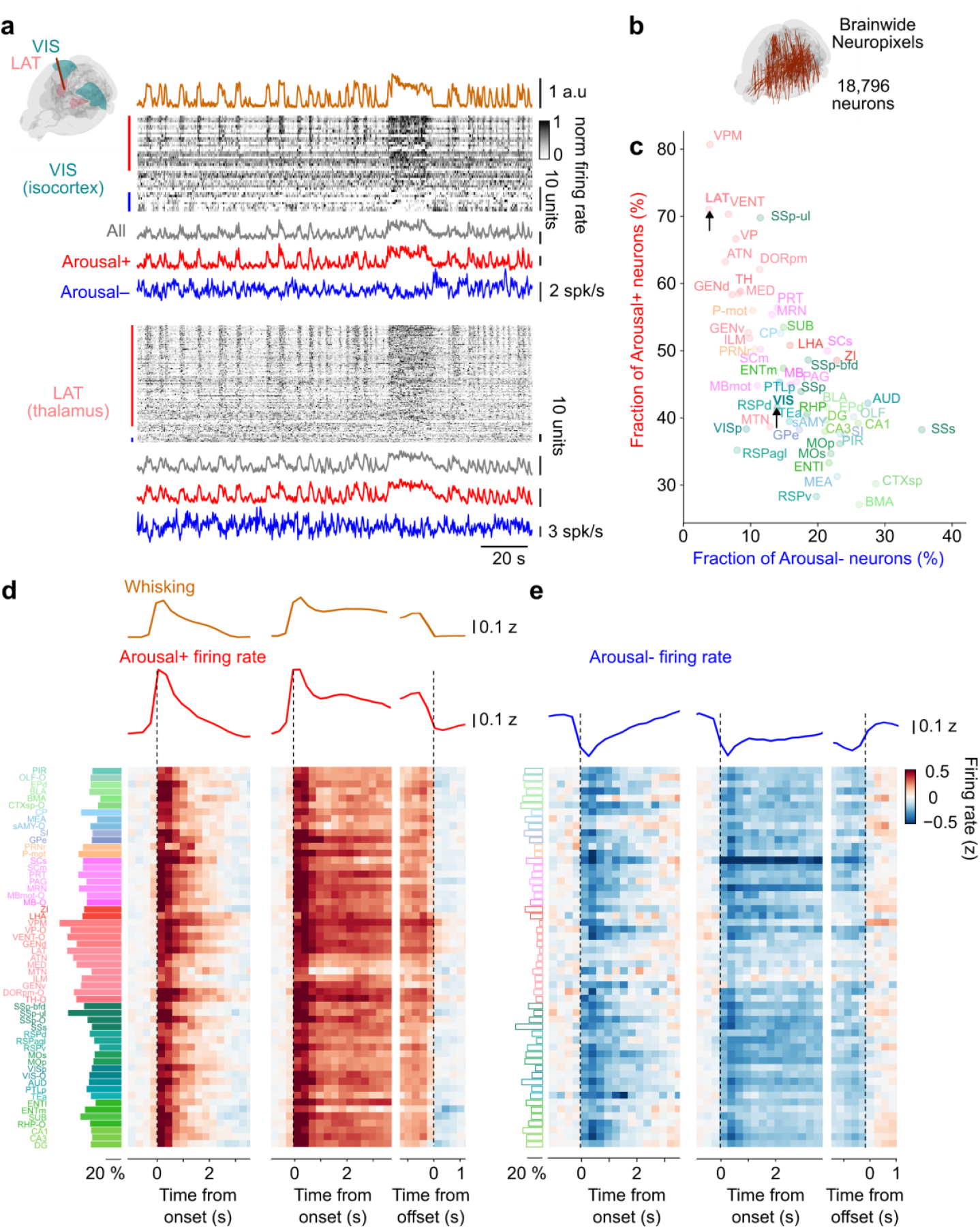
The two arousal-defined neural populations are present throughout the brain. **a.** Example Neuropixels recording with neurons in visual cortex (VIS, *top*) and lateral group of the dorsal thalamus (LAT, *bottom*) from Ref. ^40^, showing the firing rate of all recorded neurons for each region, ordered based on their correlation with whisking (*gold*). The firing rate for each neuron is divided by the 95^th^ percentile of firing rate across time. Bars indicate Arousal+ neurons (*red*, correlation with whisking > 0.05), and Arousal– neurons (*blue*, correlation with whisking < −0.05). *Bottom*: Average firing rate for all (*grey*), Arousal+ (*red*) and Arousal– neurons (*blue*). The firing rate of each neuron is z-scored before averaging. Arousal+ and Arousal– neurons are identified on half of the data, and activity is shown for a snippet of the other half. **b**. A map with all the Neuropixels probe insertions that we included for analysis (18,791 neurons across the brain^40^). **c**. Comparison of the fraction of Arousal+ and Arousal– neurons across regions. Arrows indicate the example regions shown in **a. d**. *Left*: Proportion of Arousal+ neurons across regions. *Right*: Average firing rate during whisking events for Arousal+ neurons within each region. Activity is aligned to the onset of brief bouts (*left*), the onset (*middle*) or the offset (*right*) of long bouts. The average activity of Arousal+ neurons across regions peaked at the time of whisking onset (0 s offset, *dashed line*). **e**. Same, for Arousal– neurons. The average activity of Arousal– neurons across regions hit its lowest mark after whisking onset (*dashed line*).

Arousal+ and Arousal– neurons were present throughout all brain regions. As in the simultaneous fUSI-Neuropixels measurements (Figure 3), in the visual cortex, fluctuations in arousal drove two opposing populations of neurons^38^, increasing activity in Arousal+ neurons and decreasing it in Arousal– neurons (Figure 5a). Similar results were seen in the whole brain (Figure 5b, c): averaged across regions, about half of the neurons (47±12 %, s.d. across 55 regions) correlated positively with whisking, and a smaller but substantial fraction (16±7%) correlated negatively with whisking. The relative abundance of Arousal+ and Arousal– neurons varied between regions (Figure 5c). For instance, Arousal+ neurons were more abundant in thalamus (61±10 %, s.d. across 12 thalamic regions, e.g. Figure 5a) and Arousal– neurons were more abundant in auditory cortex (Au, 27% of neurons) and secondary somatosensory cortex (SSs, 35% of neurons).

These opponent neural populations both showed monophasic responses to arousal events. In each brain region, the firing of Arousal+ and Arousal– neurons followed a roughly similar timecourse, with opposite signs (Figure 5d, e). Their contributions thus tended to cancel out in the average firing rate (Figure S12), particularly in regions where Arousal– neurons were more frequent, such as hippocampus. The two populations, however, showed subtly distinct dynamics: the activity of Arousal+ neurons peaked 0.11±0.02 s (s.e. across 55 regions) after whisking onset (Figure 5d). The activity of Arousal– neurons peaked 0.78±0.26 s after whisking onset (Figure 5e), significantly later than Arousal+ neurons (p = 8e-6, Wilcoxon signed-rank test across regions).

The distinction between Arousal+ and Arousal– neurons was robust and stable across contexts and time. Up to now, we had based this distinction on the correlation between neural activity and whisking measured in a single recording session in passive mice. To assess whether this distinction is robust to behavioural context, we analysed the responses of the same neurons while mice performed a visuomotor task^40^. To assess whether the distinction might vary across days, moreover, we implanted a chronic Neuropixels probe^60^ and tracked individual neurons^61^ for 6 or more days. In both cases, the results revealed remarkable stability: correlation with whisking was largely the same whether the mice were passive or performing a task (Figure S13a, b) and was largely constant across days (Figure S13c, d).

The combined model based on the firing rate of Arousal+ and Arousal– neurons provided a good account of brainwide blood volume during whisking events, much superior to the traditional model based on bulk neural activity. We used the bulk model (Figure 3) and the combined model (Figure 5) to predict whisking-related blood volume throughout the brain. For the bulk prediction, we convolved the bulk firing rate obtained with the brainwide Neuropixels recordings (Figure S12) with the HRF we had obtained for bulk firing rate in our simultaneous Neuropixels and fUSI recordings (Figure 2e, Figure S8, averaged across all measurements in visual cortex and hippocampus). We followed the same procedure to obtain combined predictions from of Arousal+ and Arousal– populations for each brain region, this time applying the filters we had found for Arousal+ and Arousal– neurons (Figure 3e) to the firing rate of these populations in each region (Figure 5d, e). We scaled the contributions of Arousal+ and Arousal– populations based on the relative sizes of these populations in each region, and we summed them to obtain a combined prediction of whisking-related changes in brainwide blood volume. The bulk prediction completely missed the biphasic dynamics and the delays of blood volume following whisking events(Figure 6a) whereas the combined prediction well recapitulated these characteristics (Figure 6b). The combined prediction thus largely outperformed the bulk prediction, leading to a smaller error in most brain regions (Figure 6c).

**Figure 6.**
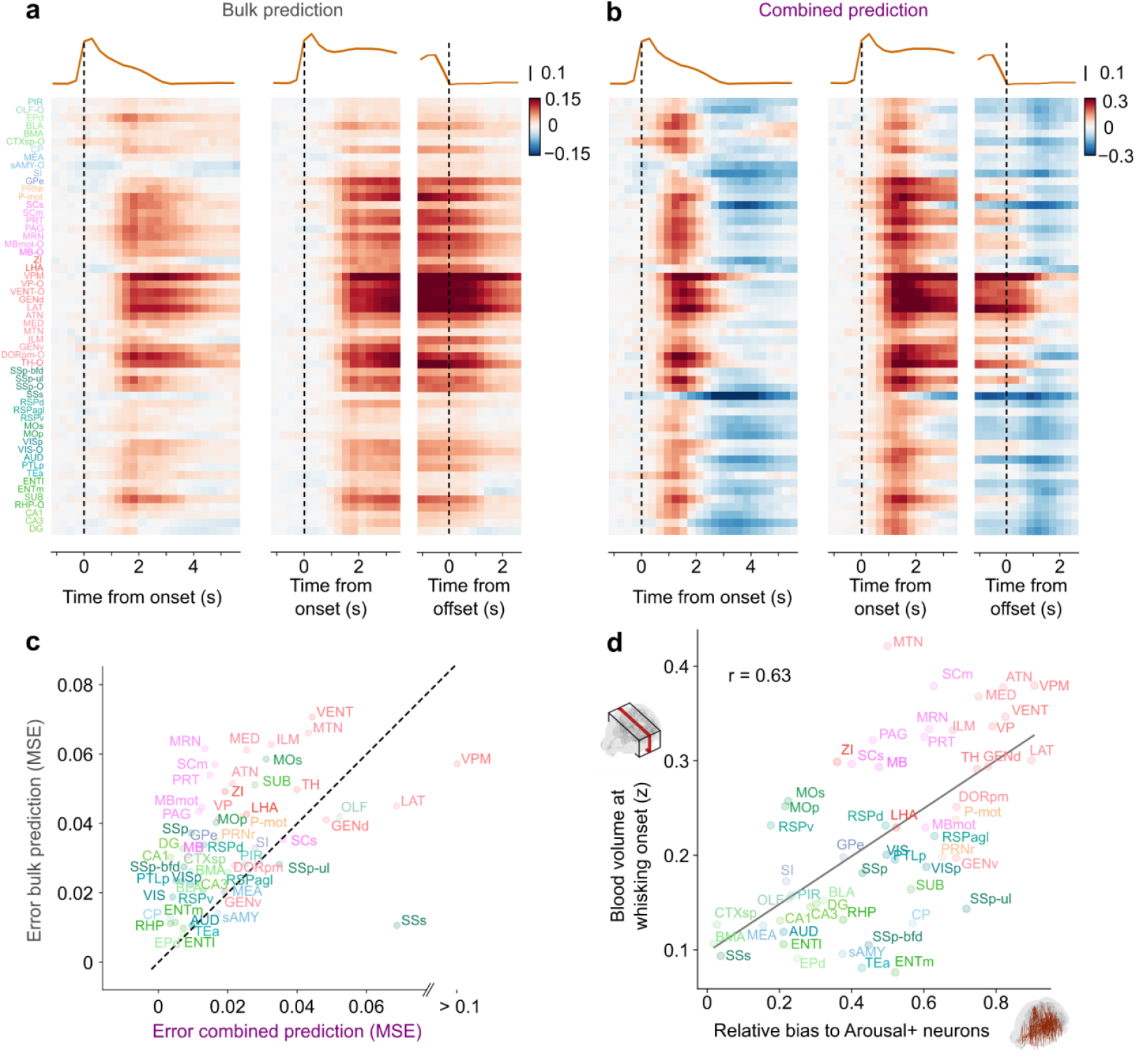
The two arousal-based populations better predict brainwide blood volume around arousal events. **a.** Bulk prediction of whisking-evoked changes in blood volume, obtained by convolving whisking-related bulk firing rate (as shown in Figure S12) with the filter from Figure 2e, averaged between visual cortex and hippocampus (Figure S8). **b**. Combined prediction of whisking-related blood volume from Arousal+ neurons and Arousal– neurons, obtained by convolving the firing rate of the two populations (as shown in Figure 5d, e) with their respective filters from Figure 3e, averaged between visual cortex and hippocampus (Figure S8) and summing their contributions scaled by the proportion of Arousal+ and Arousal– neurons in each region. **c**. Mean squared error (MSE) between actual and predicted whisking-related blood volume for the bulk prediction, plotted against the MSE for the combined prediction. The MSE is computed across time for the brief and longer whisking bouts (0–4.5 s window). Each dot is a brain region, accompanied by its Allen acronym. **d**. Average blood volume (from the brainwide fUSI recordings) 1.2-J. s after whisking onset plotted against the relative bias to Arousal+ vs. Arousal– neurons (computed as the difference in the number of Arousal+ and Arousal– neurons over their sum (from the brainwide Neuropixels recordings). Each dot is a brain region, accompanied by its Allen acronym. The linear fit (*black line*) highlights the strong Pearson correlation (r = 0.63).

In addition to yielding superior estimates of blood volume across regions, the combined model provided a rational basis for the differences in blood volume seen across regions during whisking events. The blood volume measured in each region after whisking onset could to some extent be predicted by the bulk firing rate increase measured in that region (r = 0.44 across 55 brain regions, p = 0.0008). However, the bulk firing rate increases underestimated the blood volume changes in some midbrain regions and overestimated them in some cortical regions (Figure S12). A better explanation of the blood volume across regions was the relative proportion of Arousal+ vs. Arousal– neurons in those regions, with more accurate estimates of blood volume changes for many brain regions, especially in the midbrain and cortex (r = 0.63 across regions, p = 3.3e-7, Figure 6d). The two predictors — bulk firing rate after whisking onset and abundance of Arousal+ neurons — correlate with each other, but the second one was superior in predicting blood changes across the brain. Indeed, incorporating both factors yielded predictions that were significantly better than those based on bulk firing rate alone, and not statistically different from those based solely on the relative abundance of Arousal+ neurons (Partial F-test, N = 55, p = 0.44).

## Discussion

By combining three datasets acquired in different mice with different techniques, we obtained high-quality predictions of fluctuations in blood volume across the mouse brain. Large-scale fUSI measurements revealed brainwide fluctuations in blood volume associated with whisking. Simultaneous fUSI and Neuropixels recordings revealed that neurons that increase vs. decrease activity with whisking (Arousal+ and Arousal–) have distinct, positive hemodynamic response functions (HRFs). Brainwide Neuropixels recordings revealed that these two populations of neurons coexist in the whole brain. Filtering their firing rates with their HRFs and combining the results provided good accounts of brainwide changes in blood volume with arousal, outperforming the traditional model based on bulk firing rate. Thus, neurovascular coupling appears to reflect not the bulk firing rate, but rather the firing rates of two brainwide neural populations with opposite relation to arousal.

This finding extends, without contradicting them, previous studies of neurovascular coupling. Indeed, most studies could not have observed the two opposing populations, because they obtained a single measure of bulk neuronal activity, such as the local field potential or the overall firing of a local population. A study^35^ that distinguished the vascular correlates of different neurons, indeed, found results that are consistent with ours, revealing neurons that had opposite correlations with the local vascular signal and opposite correlations with activity in brainstem regions, which might be related to arousal.

Brainwide hemodynamic fluctuations have been observed in a variety of species, from mice to humans, and are a prominent feature of all brain states, from sleep to task engagement. In rodents, fluctuations in arousal, engagement, and movement are associated with fluctuations in neural activity in multiple brain regions^38,40,49,62–65^ and with concomitant global hemodynamic signal across the cortex^12,47^ and in the rest of the brain^66^. The highly correlated fluctuations in neural activity and global hemodynamic signals^10,67,68^ may depend on arousal-related forebrain nuclei^69^ and on serotonin neurons in the Dorsal Raphe Nucleus^70^. Similarly, in non-human primates fluctuations in arousal are associated with brainwide fluctuations in fMRI BOLD signals across brain states^71^, which are highly correlated with neural activity^21,35^ and likely depend on acetylcholine released from Nucleus Basalis^72^. Similar brainwide fluctuations in hemodynamic activity are seen in humans, in the global fMRI signal. This global signal correlates with vigilance measured by EEG^73^, and with arousal measured by fluctuations in pupil dilation^74–76^, and in associated physiological signatures such as respiration or heart rate^77–79^. Such brainwide hemodynamic activations are seen not only in wakefulness but also during transitions from NREM to arousal^80^ and during NREM sleep ^78,81^.

Our results in mice indicate that a substantial part of fluctuations in blood volume throughout the brain are predictable by a measure of arousal, which we based on whisking. Arousal has multiple correlates, including pupil dilations and locomotion^12,51–54^. We found that whisking was a more effective predictor than locomotion, because locomotion was always preceded and accompanied by strong whisking whereas whisking can occur without locomotion. Though its dynamics are slower, the pupil may have provided a similarly good predictor^47^. However, our results are limited to the whisking that is observed during head fixation: we do not know if they would extend to the whisking observed during free exploration of an environment.

Fluctuations in blood volume, nonetheless, were better explained by the activity and relative abundance of Arousal+ and Arousal– neurons, which varied substantially across regions. This variation may explain why some cortical regions are overall more positively vs. negatively correlated with locomotion^37,64,82,83^. For instance, the abundance of Arousal– neurons in auditory cortex echoes the prevalence of neurons suppressed by locomotion in that region^84^.

These results would be hard to explain based on non-neural factors. The neural recordings revealed that whisking is associated with brainwide changes in neural activity, and that these changes are different across brain regions, with more Arousal+ neurons in some regions and more Arousal-neurons in others. This variation across regions helps explain the different effects of arousal on blood flow in those regions, without requiring factors other than neural ones. Moreover, the same model explains blood flow based on neural activity regardless of brain state, from REM sleep to active. It is hard to imagine that this result would arise from non-neural factors that specifically affect blood flow in some behavioural states. Such non-neural factors might be a concern had we proposed the opposite view – that one needs different models to relate neural activity to blood in different brain states.

Once the diverse abundance of the two populations across brain regions is considered, neurovascular coupling becomes consistent across the brain. Indeed, it is described by a single model with the same two HRFs across brain regions. This result may reconcile studies that proposed that neurovascular coupling is consistent^10,20,21^ vs. diverse^5,14,16–19,85^ across regions. Perhaps studies that found it to be consistent compared regions with similar fractions of Arousal+ and Arousal– neurons, and studies that found it to be variable compared regions with different fractions.

Our results also support the view that neurovascular coupling is consistent across brain states. Indeed, we found that the activity of the two opposing populations could predict blood volume across arousal states ranging from REM or NREM sleep to quiet or active wake. This result offers a reinterpretation of previous studies that found neurovascular coupling to differ across brain states^5,21,23,24^: these studies used bulk measures of neural activity, so they could not distinguish the state-dependent contributions of Arousal+ and Arousal– neurons.

The ability of the two opposing populations to predict blood volume may be even higher than it appears from our analyses. Indeed, our analyses relied on three datasets obtained in different mice, laboratories, and conditions. This uncontrolled variation is likely to affect mouse behaviour, making the comparison more difficult. For instance, the mice were likely quietest in the simultaneous fUSI and Neuropixels recordings, which were performed in long sessions with no task. By contrast, the brainwide Neuropixels recordings that we analysed were briefer and performed right after the mouse performed a task.

Moreover, the relationship between neural activity and blood volume may have higher dimensionality. We have distinguished two populations of neurons, Arousal+ and Arousal–, based on their correlation with whisking. There may be other ways to distinguish the activity of neurons that would lead to even better predictions. These improved approaches might identify more than two subpopulations, each with their own HRF.

A promising avenue for further research will be to clarify the relation between Arousal+ and Arousal– neurons, which are defined functionally, and the wide variety of cell types in the brain, which are defined genetically and which may play different roles in neurovascular coupling^29^. Presently, however, the literature on these roles shows little consensus. For instance, neurons expressing nNOS, and particularly Type-1-nNOS or Sst-Chodl cells, are ascribed a major role in neurovascular coupling, particularly during movement, but ablating them does not abolish neurovascular coupling^33^. Moreover, there is debate as to their relation to arousal: they are active during low-arousal and NREM^86^ (which would mark them as Arousal–), but they can also influence whisking-evoked blood flow, and whisking is prevalent in REM and active states. Perhaps different studies targeted slightly different cell types among the various Nos1-expressing inhibitory^87^ and excitatory neurons, which have diverse relationships to arousal^86,88–90^.

A class of neurons that might have an outsize effect on blood volume are those that release neuromodulators such as Norepinephrine (NE), which may affect blood volume independently of neural activity^13^. Indeed, a model has been proposed^13^ where vascular signals in the cortex result from a combination of bulk neural activity (with a positive HRF) and NE (with a negative HRF, as its effect is vasoconstrictive). Because NE correlates strongly with pupil dilation, this model resembles the one that we tried when we paired the bulk firing rate with pupil dilation (Figure S10). As expected, this model performed well in the visual cortex, where the pupil is on its own an excellent predictor of blood volume^47^, but not in hippocampus, where it did significantly worse than our combined model based on the activity of Arousal+ and Arousal– neurons.

While we cannot claim that the contributions of Arousal+ and Arousal– neurons are causal, it is not hard to imagine that they might be. Indeed, in the combined model both HRFs are positively lagged (changes in neural activity precede changes in blood volume) and positively signed (increases in neural activity correspond to increases in blood volume). Arousal+ and Arousal– populations might thus be each related to blood volume through their own distinct — but conventionally vasodilatory — neurovascular coupling mechanisms.

Alternatively, there could be an additional direct link between the factors that control the activity of those two populations and blood volume. These factors have been studied extensively in the cortex and in thalamus, and include neuromodulation^13,44,91–94^ and inhibition^37,95–99^. These factors may affect different populations of neurons differently. Our data do not distinguish between these causal and noncausal explanations. They do, however, reveal that neurovascular coupling is different in two key neural populations with opposite relation to arousal.

## Acknowledgments

We thank Charu Reddy for invaluable help with surgeries and lab management, Célian Bimbard for help with the chronic Neuropixels experiments, and Samuel Le Meur-Diebolt and Jacob Ratliff for helpful suggestions. This work was supported by the Wellcome Trust (grant 223144/Z/21/Z to MC and KDH) and the US BRAIN initiative (NIH grant U19NS123716 to the International Brain Laboratory). MC holds the GlaxoSmithKline / Fight for Sight Chair in Visual Neuroscience.

## Methods

### Brainwide fUSI measurements

Experiments were conducted in 5 C57/BL6 mice (3 male, 2 female), 13-34 weeks of age. All experimental procedures were conducted according to the UK Animals Scientific Procedures Act (1986). Experiments were performed at University College London, under a Project License released by the Home Office following appropriate ethics review.

#### Surgery

Mice were first implanted with a headplate and cranial window under surgical anaesthesia in sterile conditions. The cranial window replaced a dorsal section of the skull (∈8 mm in ML and ∈5 mm in AP) with 90 μm thick ultrasound-permeable polymethylpentene (PMP) film (TPX DX845 PMP, Goodfellow Cambridge Ltd.) attached to the skull with cyanoacrylate glue. The PMP film was then covered with Kwik-Cast (World Precision Instruments, USA), except during imaging sessions. After surgery, mice were allowed to recover for 7 days before starting habituation. Mice were then habituated to handling and head-fixation in the rig across several days, as session duration progressively increased.

#### Histology

Mice were perfused with PBS, PFA, and a gel containing TRITC–Dextran (Sigma-Aldrich) to perform blood vessel labelling. We imaged full 3D stacks of the brains in a custom-made serial section^100^ two-photon^101^ tomography microscope. Images were acquired using ScanImage (Vidrio Technologies) and the hardware coordinated using BakingTray. We extracted the signal in green and red channels, for autofluorescence and labelled blood vessels.

#### fUSI recording sessions

Recordings were performed in 5 mice across different days for up to 10 weeks. On a given day, we sequentially recorded 2-3 sessions in different coronal planes, lasting ~45 min each. The final dataset consisted of 138 such sessions for a total of 101 hours of recordings, ranging from 17 to 24 hours per mouse. Because each plane did not contain all regions, the exact number of recording sessions for each brain region differed (Md by animal by region: 11±7.7 s.d., see Figure S1 for all numbers).

In each recording session, we head-fixed the mice by securing the headplate to a post placed 10 cm from three video displays (Adafruit, LP097QX1, 60 Hz refresh rate) arranged at right angles to span 270 deg in azimuth and ∈70 deg in elevation. The screens were uniform grey or black. In some sessions (13 out of 138), the screens switched from grey to black, or from black to grey halfway through the recording session. The mice were placed on a running wheel. Four infrared cameras and the wheel rotary encoder were used to monitor their spontaneous behaviour.

We covered the PMP film with ultrasound gel and positioned the ultrasound transducer above it (128-element linear array, 100 μm pitch, 8 mm focal length, 15 MHz central frequency, model L22-Xtech, Vermon, France). Doppler signals from the transducer were acquired using an ultrasound system (Vantage 128, Verasonics, USA) controlled by a custom MATLAB-based user interface (Alan Urban Consulting) recording continuously at 500 Hz.

#### fUSI preprocessing and denoising

Coherent tissue motion was removed by performing SVD on the complex images (IQ data) in overlapping 600 ms windows (300 ms overlap) and discarding the first 50 principal components^102^. This number provided a trade-off between effectively removing tissue motion and preserving actual blood-related signal. Similar results were obtained when removing any number of components between 20 and 100 (Figure S14). Power Doppler images were then obtained by computing the average squared modulus of the filtered IQ data over non-overlapping 300 ms windows.

Compared to experiments in freely-moving animals^85,103^ and with measurements through the cranium^104^, motion artefacts are less of a concern when fUSI is conducted through a cranial window in well-habituated head-fixed mice. In these conditions the signals are strong, and the movements are small. Nonetheless, to minimize the possible impact of motion artefacts, we denoised the Power Doppler data using several steps:

1. To correct for brain deformation, we performed motion registration. We used NoRMCorre, a piecewise-rigid registration algorithm from CaImAn^105^. This step corrects large in-plane movements of the brain in which displacements exceed the voxel size (>0.1 mm), which typically occur during running. We note that off-plane displacements are unlikely to affect the signal, as signals are integrated over a wider distance in the antero-posterior axis (> 0.4 mm), much larger than the range of brain movements.
2. To correct for timepoints with artefactually high values (outliers), we ran a procedure to identify those and replaced them through linear interpolation of neighbouring frames. The detection was based on (a) identifying noisy voxels as those with kurtosis across time above 10, (b) computing the changes between successive frames of the average activity of these noisy voxels, (c) identifying time points in which this signal was greater than the mean by 3 standard deviations. This led to removing 0–3% of timepoints (see Figure S15 for the full distribution across sessions).
3. To correct for residual contamination of motion artefacts, we removed any activity that could be linearly predicted from out-of-brain voxels (mostly located in the ultrasound gel, above the surface of the brain). We identified out-of-brain voxels based on low average activity and not being assigned to any brain region. We computed SVD on out-of-brain voxel activity and used the first 5 components as regressors. We fitted linear weights to predict the full power doppler using ridge regression and subtracted this prediction from the data. We chose this number as a trade-off between capturing enough variance and not introducing noise through the regression. As shown in Figure S15, subsequent components only explained little additional variance.

Finally, we z-scored the denoised Power Doppler signal of each voxel across time.

After these corrections, motion artefacts are unlikely to contribute significantly to the large hemodynamic signals we observe with arousal events. First, the increases in fUSI signals seen at times of arousal appear with the typical ~1 s lag of blood signals. If they were motion artefacts, they would occur at zero lag. Indeed, such zero-lag effects appeared only when we relaxed our procedures for removing motion artefacts, for example by discarding less than 10 components in the SVD filtering (Figure S14). Second, many of the arousal events involve only tiny twitches of the nose, which are unlikely to cause artefacts. Consistently, most of our denoising steps do not affect these signals (Figure S15).

#### Alignment to Allen atlas and region extraction

To extract signals for different brain regions and combine data across sessions and animals, we used a custom, individual-based alignment to the Allen Common Coordinate Framework atlas (CCF, ^42^).

For each animal, we ran an anatomical scan using a motor interface. This scan consisted of average Power Doppler images for short fUSI acquisitions in slices 0.1 mm apart, covering the whole craniotomy. This served as a reference to which we aligned each individual session, as well as the histology.

We perfused each animal using a blood vessel labelling procedure and imaged the brain (see Histology), extracting signals in two colour channels: green for autofluorescence and red for labelled blood vessels. We aligned the volume to the Allen CCF atlas using the autofluorescence channel with brainreg^106–108^.

We then aligned the in-vivo fUSI anatomical scan of each animal to its in-vitro volume using the stained blood vessels. To do so, we used an affine transformation fit to minimize the distance between several manually identified matched key-points (~15-30) in both volumes. These were typically branching between big blood vessels visible in both the in-vivo, lower resolution fUSI scan, and the in-vitro, higher resolution histology volume. We visually confirmed the proper alignment between the two volumes and then transposed the Allen regions labels to our fUSI volume, so that we obtained a region label for each voxel of the scan.

To select the target regions to keep for our analyses, we used the coarser Beryl mapping^40^ and set a threshold of volume for regions to be considered (~0.8 mm^3^), as smaller regions are less likely to be identified with confidence with the resolution of fUSI. We merged regions smaller than this threshold to their parent region and reiterated the procedure until all regions reached the threshold. This procedure resulted in the co-existence of regions at various stages of hierarchy, for example VPM, VP and VENT. To clarify that these excluded some regions, we used the -O to indicate “other”, in VP-O and VENT-O. A full list of region names and acronyms is shown in Table S1.

fUSI off-plane (AP) spatial resolution depends on depth, but is at least 0.4 mm. To ensure that the signal in each voxel reflected a single brain region of interest, we computed its mode region label across the AP axis in a 0.4 mm range. We only assigned a region to a voxel if the mode corresponded to more than 2/3 of the values. Thus, voxels in zones of transitions between different regions were not assigned to any region.

For each recording session, we then aligned the slice to our reference anatomical scan in two steps: (a) manual assignment to the closest AP slice in the reference scan, (b) cross-correlation in space with the chosen slice of the reference scan. We then used the region labels from the aligned reference scan.

We discarded voxels with very little signal due to shadowing, for e.g. due to bone regrowth. To do so, we applied a Sobel filter to the anatomical image, which computes an approximation of the gradient of the intensity of the image, or edges. We identified shadowed areas by patches of the image (> 30 voxels) with consistently low values of the Sobel filtered image. Indeed, in the brain, individual vessels create many edges, so that big patches of low values only occur outside of the brain, or when vessels are hidden by bone.

Once we obtained region labels for each recorded voxel, we computed the region signal for this session using the median activity across voxels. For each region, we only included sessions in which are least 20 voxels of the region were present in the slice. Region signals were extracted for each hemisphere separately and then averaged across the two hemispheres. For further analysis, we considered all regions for which we recorded at least 10 sessions in total over all animals.

#### Behaviour analysis and extraction of onsets

Locomotion velocity was obtained from the rotary encoder.

Whisking measurements were obtained through Facemap^38^. Whisking was obtained by taking the first component of a SVD decomposition of motion energy over the whisker pad. This was done on either one or two cameras (corresponding to both sides of the mouse face). If the signal was available for two cameras, we used the average across both cameras. To have a comparable signal across sessions, we ensured that higher values corresponded to higher motion, and normalized the signal between 0 and 1, based on the 1^st^ percentile and the maximum.

All behavioural metrics were binned in the same 300 ms bins as the fUSI data.

Whisking bouts were detected when the signal crossed a given threshold (0.15) and lasted until the signal reached another threshold (0.05) under the condition that the average whisking in the next 4 s remained below this threshold (this is to exclude very short interruptions in longer whisking bouts and allow better separation of whisking bouts).

Brief bouts were defined as bouts with duration between 1.3 and 3.5 s, while longer bouts lasted longer than 3.5 s. Whisking bouts were considered accompanied by locomotion when wheel velocity above 1 cm/s was detected at any point in the 6 seconds following whisking onset.

#### Whisking-evoked activity

To compute whisking-evoked activity for all regions, we first computed results for each mouse and then averaged across mice. For each region and each mouse, we aligned data to the detected onsets (or offsets) and averaged across all recorded whisking events of a given category (e.g. short or longer bouts, with or without locomotion). Before averaging, for each bout we excluded times when another, following whisking bout had started. We then averaged results across animals, only keeping regions with more than 10 recording sessions in total. Finally, we recentred by the baseline activity (0.5–2 s before whisking onset). Because not all regions were recorded in all sessions, whisking-evoked activity for each region reflects a variable number of sessions, and thus of whisking events. We report the number of sessions and events considered for each region and animal in Figure S1.

#### Statistical testing

To assess the statistical significance of the changes in blood volume associated with whisking events (Figure 1g–i, Figure S4), we used a permutation test. For each animal, we randomly shuffled the mapping between sessions and whisking and locomotion velocity, so that we aligned blood volume to the whisking events that occurred in another session. We repeated this procedure 100 times, thus obtaining a null distribution of event-evoked changes in blood volume. For each region and timepoint, we compared the ‘true’ value of blood volume change to the null distribution and defined the p-value as the proportion of samples from the null distribution above (or below) the ‘true’ value. We corrected for multiple comparisons by using the false discovery rate. We report the proportion of regions that show significant (p-value < 0.05) increases or decreases in activity for each timepoint relative to whisking events (Figure 1i). For select timepoints (0.9 s and 3.0 s), we mark the significant regions with an asterisk (Figure 1g, h).

#### Predictions from whisking

To predict blood volume from whisking, we used cross-validated ridge regression. The lambda hyperparameter used for regularization was fitted through nested cross-validation. A feature matrix was constructed using lagged versions of the whisking trace, with 25 lags ranging from −1.8 to 5.7 s in 0.3 s intervals, where positive lags mean that whisking precedes blood volume. This matrix was used to predict activity in each brain region, for each session. For each session, we allowed for a different non-linear scaling of whisking (the same across all regions). We ran predictions from whisking raised to the power of different values (from 0.1 to 1.2 with steps of 0.1) and selected the value providing the best prediction accuracy (averaged across regions). The median exponent across sessions was 0.4, so that this procedure gave more relative weight to small whisking events relative to the strong whisking associated for example with locomotion (Figure S2).

### Combined fUSI and Neuropixels recordings

#### Imaging and recordings

For combined fUSI and neuronal recordings, we used our published dataset^10^ which combines Neuropixels recordings in visual cortex and hippocampus with fUSI in 5 C57/BL6 mice (4 male, 1 female), 9-12 weeks of age (see Ref^10^ for details). The dataset comprises 10 independent recording sessions, each corresponding to an acute Neuropixels insertion, and thus sampling different neurons. In each recording session, 3 to 5 different fUSI planes were sampled sequentially, each time repeating the same protocol including spontaneous activity (grey screen) and visual stimulation (flickering checkerboards).

Power Doppler was recomputed from IQ data to match the pre-processing steps explained above. Blood volume was averaged within two regions of interest, corresponding to voxels immediately around the probe track and in either visual cortex or hippocampus (~50 voxels in each region). In 5/10 recordings, two probes were inserted bilaterally. In these recordings, for each region we averaged the corresponding fUSI signal across the two hemispheres and considered neurons across the two probes. Units selected were ‘good’ and ‘multi-unit’ clusters from Kilosort as in Ref^10^. Blood volume, behavioural metrics and single-unit firing rate were all binned in 300 ms bins.

#### Behaviour analysis

Mice were not on a running wheel so that there is no locomotion in this dataset. An infrared camera over the right side of their face was used to monitor behaviour. Whisking was measured as the motion energy in a region of interest focused on the whisker pad, binned in 300 ms windows, and normalized as previously (between 1^st^ percentile and maximum). We also tracked the pupil using DeepLabCut^109^ and extracted pupil size (diameter), eye position and eye movements based on markers on the top, bottom, left and right extremes of the pupil.

#### Definition of Arousal+ and Arousal– populations

We computed the Pearson correlation with whisking in two splits of the session. We defined neurons as ‘Arousal+’ if this correlation was above 0.05 on both splits of the data, and ‘Arousal–’ if it was below −0.05 on both splits. We first z-scored the firing rate of each neuron throughout the session and then averaged these across neurons of each group (Arousal+, Arousal–, or all).

To relate Arousal+ and Arousal– neurons to previously described “positive-delay” and “negative-delay” neurons ^49,50^, we aligned the z-scored average firing rate of all neurons to global events, which we operationalized as corresponding to whisking events (Figure 3, Figure S5). We computed the centre of mass of each neuron relative to these events by summing the dot product of each neuron’s whisking-related activity (normalized between 0 and 1) and the corresponding lags and dividing it by the sum of its normalized whisking-related activity. We then computed principal component analysis (PCA) on the z-scored (neurons * lags) matrix and looked at the first component (PC1), which mapped onto the correlation with whisking and thus well segregated Arousal+ and Arousal– neurons.

#### Predictions from firing rate

We fit filters to predict blood volume from the average firing rate(s) of either all neurons, or Arousal+ and Arousal– neurons using linear regression with Lasso regularization. We did so separately for each region and each fUSI plane within a recording session, using all corresponding timepoints (not only whisking bouts). We used lags ranging from −2.1 s to 6.6 s in 0.3 s intervals (30 values in total), where positive lags mean that firing rate precedes blood volume.

Since the resolution of our HRFs is 3.33 Hz, we could only characterize variations in blood volume that are slower than 1.67 Hz. This is not a limitation, because most of the power of changes in blood volume occur at lower frequencies. In our experiments, for instance, 84% of the power was below 1 Hz.

Cross-validation was done using folds of ~8 min, with nested cross-validation for the regularization hyperparameter. In each fold, a chunk of ~8 min of data was left out, and the rest of the session (including different states, not only whisking epochs) was used to fit the HRF weights. These weights were then used to predict the left-out data. This procedure was repeated to obtain cross-validated predictions for the whole session. Filters were then computed by averaging the weights obtained across cross-validation folds. The definition of Arousal+ and Arousal– neurons was done on the whole recording session and not included in the cross-validation loop. We report our results by first averaging the metrics (e.g. coherence, filters) obtained in different fUSI planes within the same recording session (so, with the same neurons). We then report the mean, standard errors of the mean and statistical testing across the different, independent recording sessions (n = 10 for each region).

As shown in Figure S10, we also tested alternative models by fitting filters to predict blood volume from: (i) Two random populations of neurons; (ii) Whisking; (iii) Arousal+ or Arousal– neurons alone; (iv) Bulk firing rate and pupil size.

For the first analysis (two random populations), we matched the size of the populations with that of the Arousal+ and Arousal– neurons. This resulted in a prediction with a similar number of parameters as the combined model, thus controlling for the increased number of degrees of freedom of the combined vs. bulk model. We estimated the degrees of freedom for the different HRF models by counting the number of non-zero weights out of the 30 possible (from −2.1 to 6.6 s in 0.3 s steps). We did so for each cross-validation fold, and then averaged across folds in each session, and then across recording sessions. We find an average of 22.6 degrees of freedom for the bulk model, 40.7 for the combined model, and 41.3 for the random combined model. The latter yielded predictions even worse than the bulk model, so the increased degrees of freedom alone cannot account for the superior performance of the combined model.

Finally, we looked at predictions obtained by applying biphasic filters to the bulk firing rate (Figure S11). We tested four such filters, obtained by taking the weighted difference between the filter we obtained for Arousal+ neurons (weight *k*) and the filter we obtained for Arousal– neurons (weight *1-k*), with *k* taking the values 0, 0.33, 0.66, and 1. This was done for each recording session using the filters obtained for this specific session.

#### Fit evaluation

To compare whisking-evoked activity for different types of signals (firing rate, actual and predicted blood volume), we identified whisking events as previously, aligned the signal to these events and averaged across them. All traces were z-scored throughout the whole recording session prior to this alignment.

To evaluate fit quality, we computed the coherence throughout the session between actual and (cross-validated) predicted blood volume, which is like correlation in the frequency domain. Specifically, we estimated the magnitude squared coherence estimate using Welch’s method with 15 s windows and 7.5 s overlap between windows. We also computed the coherence separately for different parts of the session (Figure S9): spontaneous activity (grey screen) vs. visual stimulation (flickering checkerboards).

To compare different predictions, we ran paired t-tests between the average coherence between actual and predicted blood volume (for all frequency bands below 0.7 Hz) for pairs of predictions (n = 10 independent recording sessions per region).

#### Classification of arousal states

We used pupil size and eye movements to define four putative states: rapid-eye-movement (REM) sleep, non-rapid-eye-movement (NREM) sleep, quiet wake and active wake. We base this classification on previous work showing that pupil size alone well predicts the different sleep and wake states^12^. For each recording session, we plotted eye movements against pupil size and manually identified two critical points. Starting from intermediate pupil size, where eye movements are scarce, we identified the points at which the levels of eye movements start increasing, (i) when going to smaller pupil size – likely reflecting REM; (ii) when going to higher pupil size – likely reflecting active wake. We then used the mid-point between these thresholds as an arbitrary transition from NREM to quiet wake.

For each session, we defined 3 s intervals, measured the median of different metrics in each interval and binned them based on the median pupil size within the corresponding interval. We defined an arousal index which corresponds to the interpolated pupil size for each session, so that the previously determined thresholds correspond to 0 (REM transition) and 1 (active wake transition). We then averaged metrics across sessions for similar values of the arousal index. This allowed us to combine different sessions, where slightly different camera angles prevented us from directly comparing the absolute values of pupil size and eye movements. We validated our definition by looking at other behavioural metrics as a function of pupil size (eye position, whisking).

To evaluate prediction accuracy across states, we cut each recording session into 10 s intervals and assigned a state to each using the same method. We then computed the Pearson correlation between actual and predicted blood volume within each of these intervals and took the median correlation across all intervals corresponding to each state to obtain a prediction accuracy by state for each recording session. We then compared prediction accuracy across models in each state with a t-test across sessions (n = 10 independent recording sessions per region).

As a control, we also fit different models for each state separately. In each recording session, for each state we fit different filters to predict blood volume for firing rate using only timepoints belonging to intervals assigned to this state. We reconstructed a single prediction for the whole session by combining the cross-validated predictions obtained for each state with these different filters.

### Longitudinal recordings

To verify the stability of the correlation with whisking across days, we implanted chronic Neuropixels^60^ in two mice (one in visual cortex, one in somatosensory cortex) and tracked neurons across days using UnitMatch^61^. We then compared the correlation with whisking across recording sessions for pairs of matched neurons, depending on how many days separated them. During the recordings, mice were allowed to run on the wheel and were either facing a dark screen or presented with natural images. We excluded sessions in which the mouse ran more than 90% of the time.

### Brainwide Neuropixels recordings

We used Neuropixels recordings in the mouse brain performed in our consortium of laboratories as detailed elsewhere^40^. In the present analysis we included all sessions in which both video recordings and passive periods were available (n = 324 sessions). We focused our analysis on good units (N = 18,791 neurons) recorded in the passive periods, which occurred after the mice had performed the task.

In these experiments, mice were head-fixed onto a fixed platform (not a running wheel), so there was no locomotion. Whisking was measured as the motion energy in a region of interest focused on the whisker pad, averaged across 1 to 2 cameras, binned in 300 ms windows, and normalized as previously (between 1^st^ percentile and maximum).

We mapped neurons recorded across different animals and sessions to similar brain regions as measured in fUSI. To do so, we started with each neuron’s label in the fine-grained Allen CCF labels. We then sorted the target brain regions by hierarchical level, starting with the lowest. For a given region, we included all neurons belonging to descendants of a region except if they had been already included in another child region. For example, VENT could contain neurons belonging to the subregions of VL, VAL, but not VP, since the latter was also a target region of interest. This procedure aimed to mimic what had been done for fUSI voxels, as well as ensuring that each neuron only featured in one target region. We excluded from further analysis regions containing less than 50 neurons for which spikes were detected during the passive periods.

Spikes of each neuron were binned in the same 300 ms bins as whisking. For each neuron, we z-scored the firing rate of each neuron across time within its recording session, detected whisking events as previously and aligned this signal to the whisking events. For each type of event (brief or longer bouts), we only included neurons for which at least 5 bouts were detected.

We computed the Pearson correlation with whisking in two splits of the session. We defined neurons as ‘Arousal+’ if this correlation was above 0.05, and ‘Arousal–’ if below −0.05. To count the number of reliable Arousal+ and Arousal– neurons in each region, we only considered neurons meeting the criterion in two splits of the session. We also computed an index measuring the relative bias to Arousal+ neurons (Figure 6d) in each region by taking the difference between the number of Arousal+ and Arousal– neurons over their sum.

To establish the stability of our population’s definition across behavioural contexts, we compared for each neuron the correlation with whisking computed on the passive part of the session vs. the active part, i.e. when the mice performed the IBL task. We then computed the correlation of these values across neurons for each region (Figure S13).

To look at whisking-related firing rate, we classified neurons based on one split and computed whisking-related changes in the other split (Figure 5, Figure 6, Figure S12). All whisking-related changes are baseline-subtracted (for each region), using a window of 0.5–2 s before whisking onset.

### Predictions of brainwide whisking-related blood volume

To predict changes in blood volume (measured in the brainwide fUSI experiments) from the firing of neurons (measured in the brainwide Neuropixels recordings), we used the HRFs found from the combined fUSI-Neuropixels recordings, averaged across visual cortex and hippocampus. We assigned a neuron to Arousal+ or Arousal– groups based on half of the recording session and used its average firing rate on whisking bouts on the other half of the session. For each region, we then averaged this cross-validated firing rate across all Arousal+ or Arousal– neurons and convolved it with their respective filters. We first rescaled the Arousal+ filter by the ratio of the proportion of Arousal+ neurons in this region to the average proportion of Arousal+ neurons in the simultaneous fUSI-Neuropixels recordings (across hippocampus and visual cortex). We did the same for Arousal– filters and summed the contributions from both filters. For the predictions from bulk firing rate, we used the corresponding filter with the same scaling for all regions and used firing for the same half as for the other prediction. Finally, we applied a global scaling factor to each type of prediction (the same across all regions). This scaling factor was obtained by minimizing the mean squared error between the predicted and actual blood volume −1.2 to 5 s from whisking onset, for brief and long bouts, across all regions. We similarly computed the mean squared error for each region separately and compared those between the combined and bulk prediction (Figure 6c).

### Visualization

The 3D brain visualizations were generated using [Urchin](https://virtualbrainlab.org/urchin/installation_and_use.html).

## Supplementary materials

**Table S1:** Names and acronyms of brain regions used in this study. Each row corresponds to a brain region, with its name and acronym written in colour. Parent regions are indicated in black. Columns indicate hierarchical level as defined by the Allen Brain atlas (*left* to *right:* coarser to finer). Acronyms followed by -O (for ‘Other’) correspond to the described region excluding its child regions which are also brain regions considered in the study.

|  |  |  |  |  |  |  |  |
| --- | --- | --- | --- | --- | --- | --- | --- |
| Piriform area | CH | CTX | CTXpl | OLF | PIR |  |  |
| Cortical amygdalar area posterior part | CH | CTX | CTXpl | OLF | COA | COAp |  |
| Olfactory areas - other | CH | CTX | CTXpl | OLF-O |  |  |  |
| Endopiriform nucleus, dorsal part | CH | CTX | CTXsp | EP | EPd |  |  |
| Basolateral amygdalar nucleus | CH | CTX | CTXsp | BLA |  |  |  |
| Basomedial amygdalar nucleus | CH | CTX | CTXsp | BMA |  |  |  |
| Cortical subplate - other | CH | CTX | CTXsp-O |  |  |  |  |
| Caudoputamen | CH | CNU | STR | STRd | CP |  |  |
| Medial amygdalar nucleus | CH | CNU | STR | sAMY | MEA |  |  |
| Striatum-like amygdalar nuclei - other | CH | CNU | STR | sAMY-O |  |  |  |
| Substantia innominata | CH | CNU | PAL | PALv | SI |  |  |
| Globus pallidus, external segment | CH | CNU | PAL | PALd | GPe |  |  |
| Pallidum dorsal region - other | CH | CNU | PAL | PALd-O |  |  |  |
| Pontine reticular nucleus | BS | HB | P | P-sat | PRNr |  |  |
| Pons, motor related | BS | HB | P | P-mot |  |  |  |
| Substantia nigra, reticular part | BS | MB | MBmot | SNr |  |  |  |
| Superior colliculus, motor related | BS | MB | MBmot | SCm |  |  |  |
| Superior colliculus, sensory related | BS | MB | MBsen | SCs |  |  |  |
| Pretectal region | BS | MB | MBmot | PRT |  |  |  |
| Periaqueductal gray | BS | MB | MBmot | PAG |  |  |  |
| Midbrain reticular nucleus | BS | MB | MBmot | MRN |  |  |  |
| Midbrain motor related - other | BS | MB | MBmot-O |  |  |  |  |
| Midbrain - other | BS | MB-O |  |  |  |  |  |
| Mammillary body | BS | IB | HY | MEZ | MBO |  |  |
| Hypothalamic medial zone - other | BS | IB | HY | MEZ-O |  |  |  |
| Zona incerta | BS | IB | HY | LZ | ZI |  |  |
| Lateral hypothalamic area | BS | IB | HY | LZ | LHA |  |  |
| Hypothalamic lateral zone - other | BS | IB | HY | LZ-O |  |  |  |
| Periventricular zone | BS | IB | HY | PVZ |  |  |  |
| Periventricular region | BS | IB | HY | PVR |  |  |  |
| Ventral posteromedial nucleus of the thalamus | BS | IB | TH | DORsm | VENT | VP | VPM |
| Ventral posteromedial complex - other | BS | IB | TH | DORsm | VENT | VP-O |  |
| Ventral group of the dorsal thalamus - other | BS | IB | TH | DORsm | VENT-O |  |  |
| Geniculate group, dorsal thalamus | BS | IB | TH | DORsm | GENd |  |  |
| Thalamus, sensory-motor cortex related - other | BS | IB | TH | DORsm-O |  |  |  |
| Lateral group of the dorsal thalamus | BS | IB | TH | DORpm | LAT |  |  |
| Anterior group of the dorsal thalamus | BS | IB | TH | DORpm | ATN |  |  |
| Medial group of the dorsal thalamus | BS | IB | TH | DORpm | MED |  |  |
| Midline group of the dorsal thalamus | BS | IB | TH | DORpm | MTN |  |  |
| Intralaminar nuclei of the dorsal thalamus | BS | IB | TH | DORpm | ILM |  |  |
| Geniculate group, ventral thalamus | BS | IB | TH | DORpm | GENv |  |  |
| Epithalamus | BS | IB | TH | DORpm | EPI |  |  |
| Thalamus, polymodal cortex related - other | BS | IB | TH | DORpm-O |  |  |  |
| Thalamus - other | BS | IB | TH-O |  |  |  |  |
| Primary somatosensory area, nose | CH | CTX | CTXpl | Isocortex | SS | SSp | SSP-n |
| Primary somatosensory area, barrel field | CH | CTX | CTXpl | Isocortex | SS | SSp | SSP-bfd |
| Primary somatosensory area, lower limb | CH | CTX | CTXpl | Isocortex | SS | SSp | SSP-ll |
| Primary somatosensory area, upper limb | CH | CTX | CTXpl | Isocortex | SS | SSp | SSP-ul |
| Primary somatosensory area - other | CH | CTX | CTXpl | Isocortex | SS | SSp-O |  |
| Supplemental somatosensory area | CH | CTX | CTXpl | Isocortex | SS | SSs |  |
| Retrosplenial area, dorsal part | CH | CTX | CTXpl | Isocortex | RSP | RSPd |  |
| Retrosplenial area, lateral agranular part | CH | CTX | CTXpl | Isocortex | RSP | RSPagl |  |
| Retrosplenial area, ventral part | CH | CTX | CTXpl | Isocortex | RSP | RSPv |  |
| Secondary motor area | CH | CTX | CTXpl | Isocortex | MO | MOs |  |
| Primary motor area | CH | CTX | CTXpl | Isocortex | MO | MOp |  |
| Primary visual area | CH | CTX | CTXpl | Isocortex | VIS | VISp |  |
| Visual areas - other | CH | CTX | CTXpl | Isocortex | VIS-O |  |  |
| Auditory areas | CH | CTX | CTXpl | Isocortex | AUD |  |  |
| Posterior parietal association areas | CH | CTX | CTXpl | Isocortex | PTLp |  |  |
| Temporal association areas | CH | CTX | CTXpl | Isocortex | TEa |  |  |
| Entorhinal area, lateral part | CH | CTX | CTXpl | HPF | RHP | ENT | ENTl |
| Entorhinal area, medial part dorsal zone | CH | CTX | CTXpl | HPF | RHP | ENT | ENTm |
| Subiculum | CH | CTX | CTXpl | HPF | RHP | SUB |  |
| Retrohippocampal region - other | CH | CTX | CTXpl | HPF | RHP-O |  |  |
| Field CA1 | CH | CTX | CTXpl | HPF | HIP | CA | CA1 |
| Field CA3 | CH | CTX | CTXpl | HPF | HIP | CA | CA3 |
| Dentate gyrus | CH | CTX | CTXpl | HPF | HIP | DG |  |

**Figure S1.**
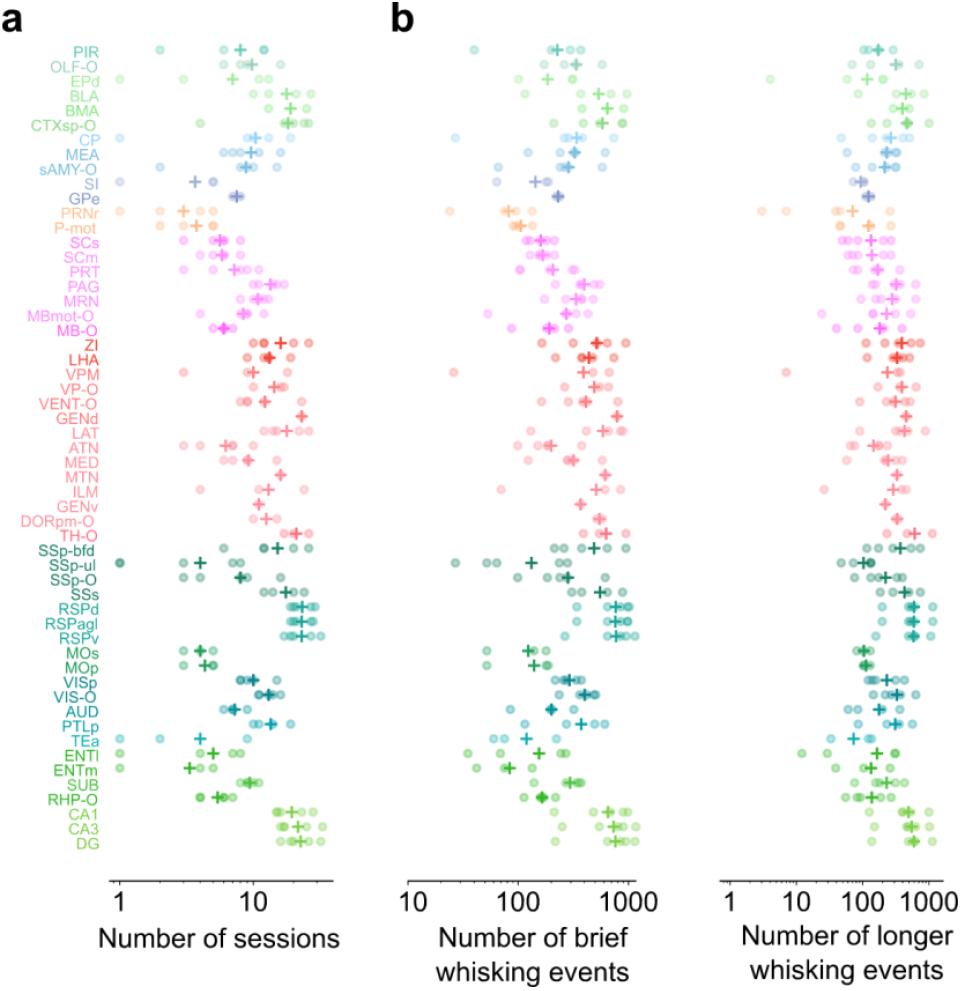
Number of recording sessions and events for each region. **a**. Number of recording sessions included for each animal (dots). Crosses indicate the average across animals. **b**. Number of brief and longer whisking events for each animal (dots). Crosses indicate the average across animals. In Figure 1g–h, whisking-evoked activity is first averaged across all events for each animal, and then across animals.

**Figure S2.**
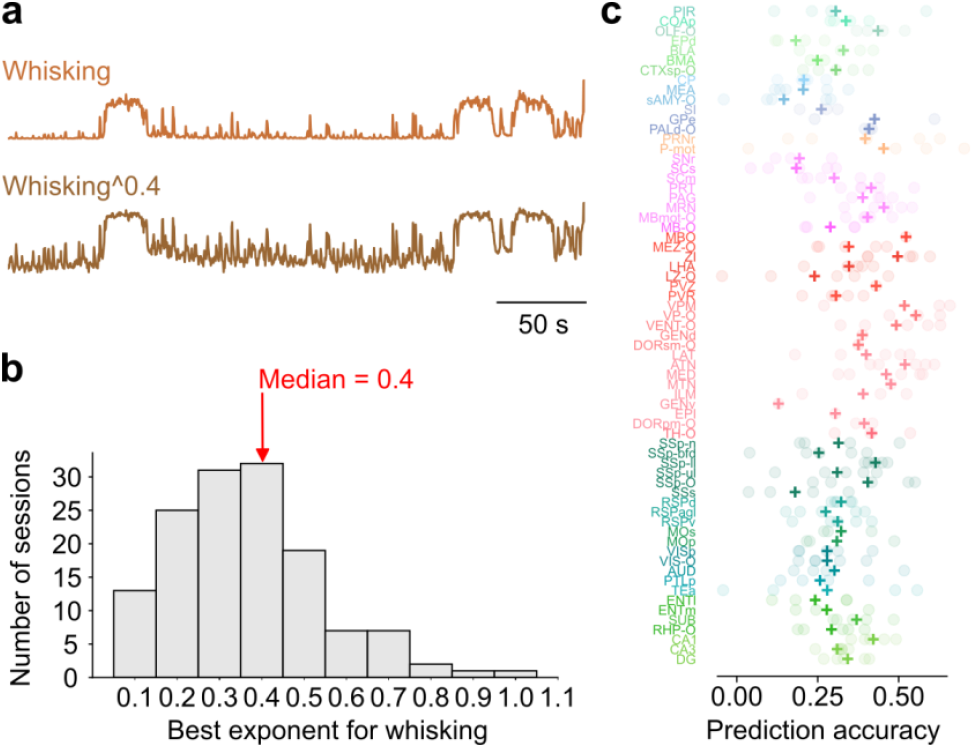
Prediction of blood volume by whisking. For each region, we used linear regression to obtain filters to predict blood volume from whisking. **a**. We use exponentiated versions of whisking, with exponents ranging from 0.1 to 1.2 in 0.1 steps. Exponents < 1, like in this example, boost small fluctuations in whisking relative to larger whisking events (which are typically associated with locomotion). For each session, we use the best exponent for that session. **b**. Histogram of best values of exponents applied to whisking (which led to highest prediction accuracy), across all sessions. **c**. Median prediction accuracy, computed as the Pearson correlation across time between actual blood volume and cross-validated predictions, averaged across sessions for each brain region. Each dot corresponds to a mouse, and the cross indicates the mean across animals.

**Figure S3.**
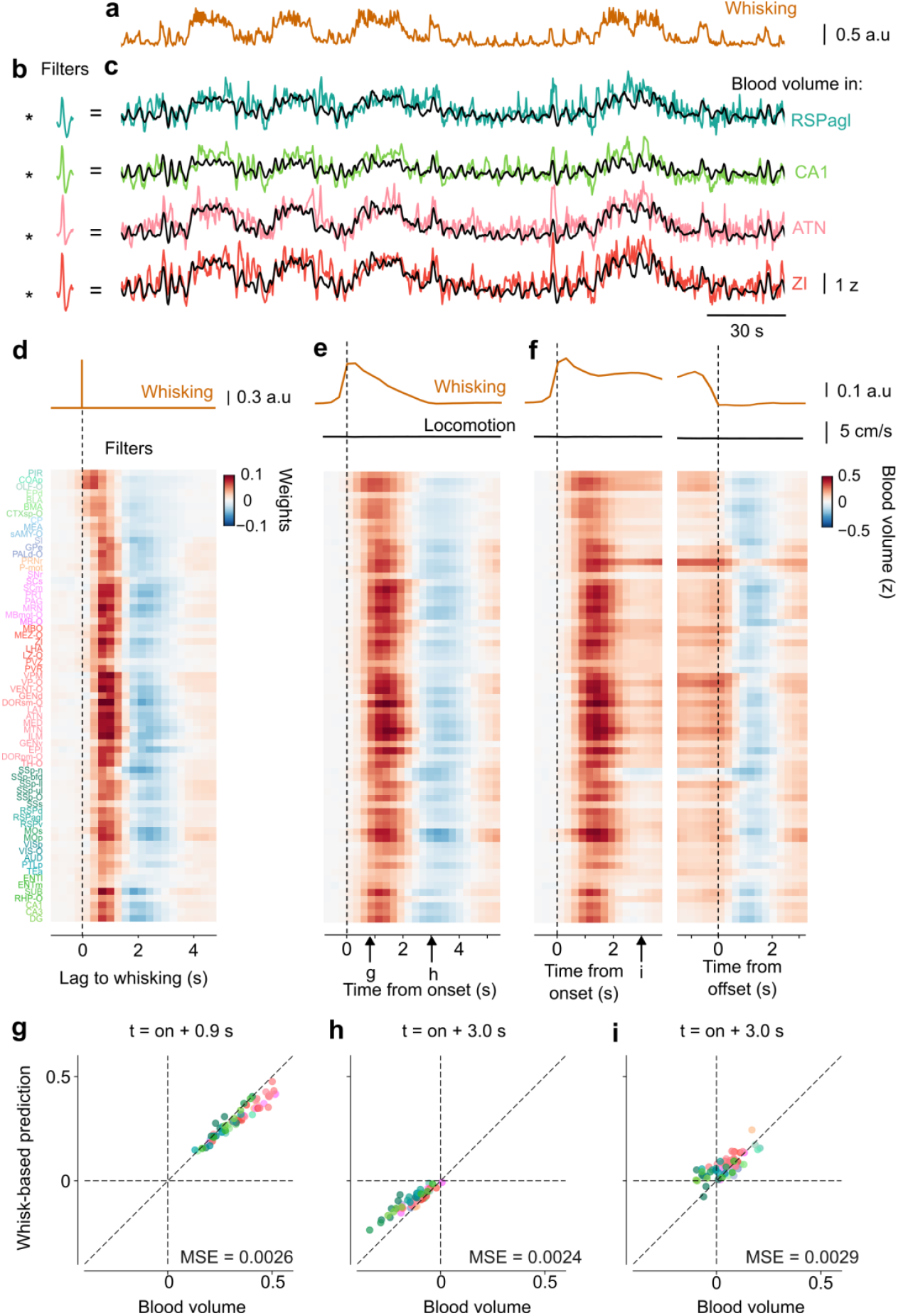
Whisking predicts brainwide blood volume throughout arousal events. **a**. For each region, we used linear regression to obtain filters that predict that region’s blood volume from whisking. These predictions were improved by applying a compressive nonlinearity to the whisking traces, which boosted small whisking events (Figure S2). **b**. Best-fitting filters to predict blood volume from whisking, in the 4 example brain regions shown in Figure 1. **c**. Blood volume measured in those 4 regions (*coloured curves*), and predictions obtained by filtering the whisking trace in a with the filters in **b** (*black curves*). **d**. Best-fitting filters obtained for all brain regions, which model changes in blood volume for a Dirac delta function of whisking. **e**–**f**. Predicted blood volume fluctuations throughout all brain regions, for brief and longer whisking bouts without locomotion. Format as in Figure 1g-h. **g**–**h**. Comparison of predicted and actual blood volume for the brief whisking events in **e**, 0.9 s (**g**) and 3.0 s (**h**) after whisking onset. Values of the mean squared error (MSE) between predicted and measured values across regions are indicated. Each dot is a brain region, coloured as in **d. i**. Same as **h**, for the longer bouts in **f**.

**Figure S4.**
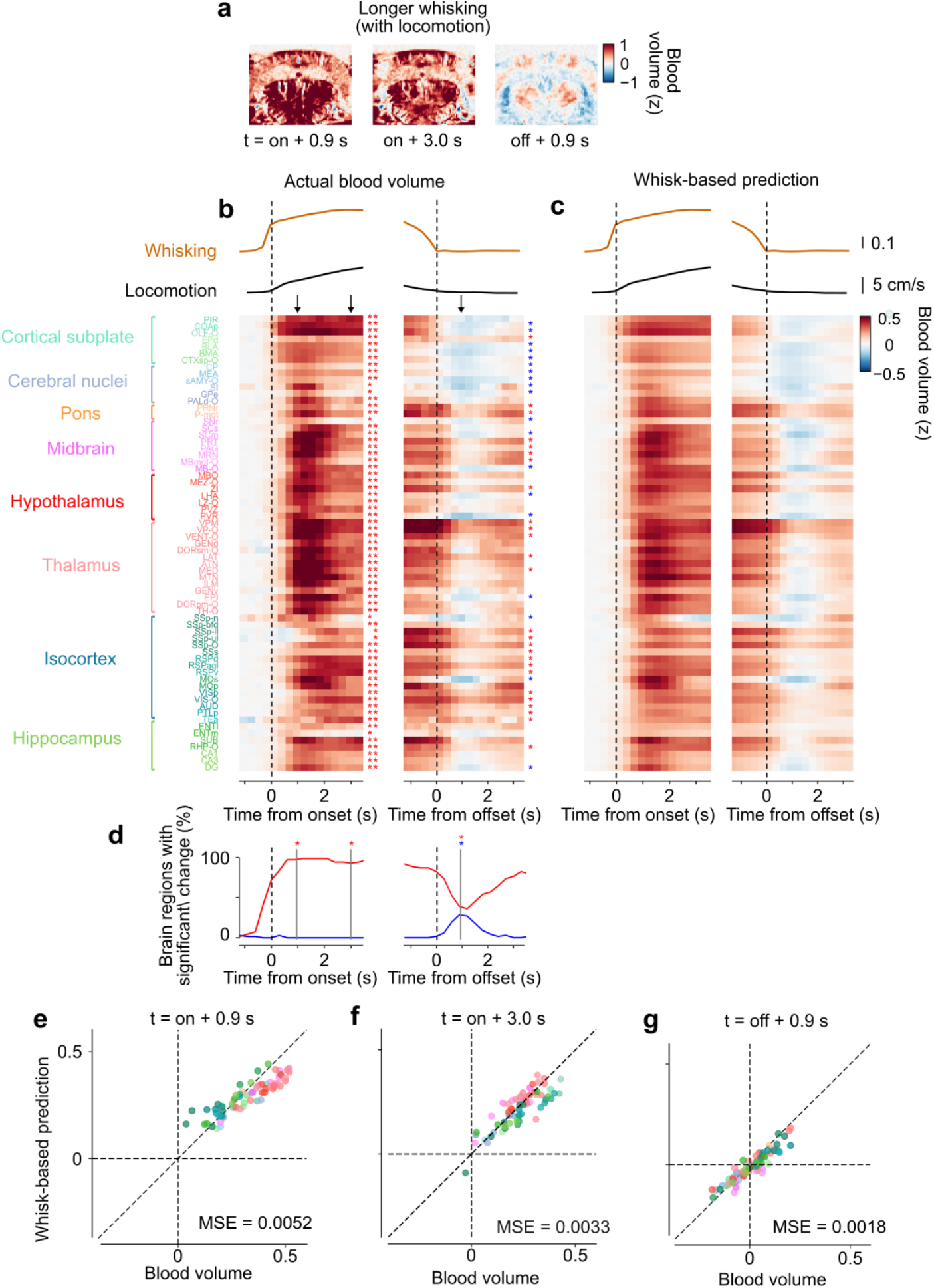
Whisking also predicts locomotion-evoked blood volume. **a**. Average blood volume for longer whisking bouts (> 3.5 s) accompanied by locomotion, 0.9 and 3.0 s after whisking onset, and 0.9 s after whisking offset, for the same example session as in Figure 1b–f. Each voxel is baseline-subtracted using a window of 0.5–2 s before whisking onset. **b**. Average changes in blood volume throughout all imaged regions during long whisking bouts accompanied by locomotion (> 3.5 s, average = 26.6 s), aligned to bout onset (*left*) and offset (*right)*. Arrows indicate the times of the images in **a** and the quantifications in **e**–**g**. Colours indicate z-scored blood volume. Results are averaged first within each animal, and then across animals (number of sessions and events indicated in Figure S1). The asterisks indicate significant changes (increases in red, decreases in blue) at times 0.9 s and 3.0 s, compared to a permutation test. **c**. Same, for predictions of blood volume from whisking (as in Figure S3). **d**. Percent regions with significant change in blood volume at each timepoint, determined using a session permutation test. Dotted lines indicate the timepoints for which significance is indicated in **b. e**–**f**. Comparison of predicted and actual blood volume 0.9 s (**e**) and 3.0 s (**f**) after the onset of long whisking bouts accompanied by locomotion. The mean squared error (MSE) between predicted and measured values across regions is indicated. Each dot is a brain region, coloured as in **b. g**. Same as **e**, 0.9 s after whisking offset.

**Figure S5.**
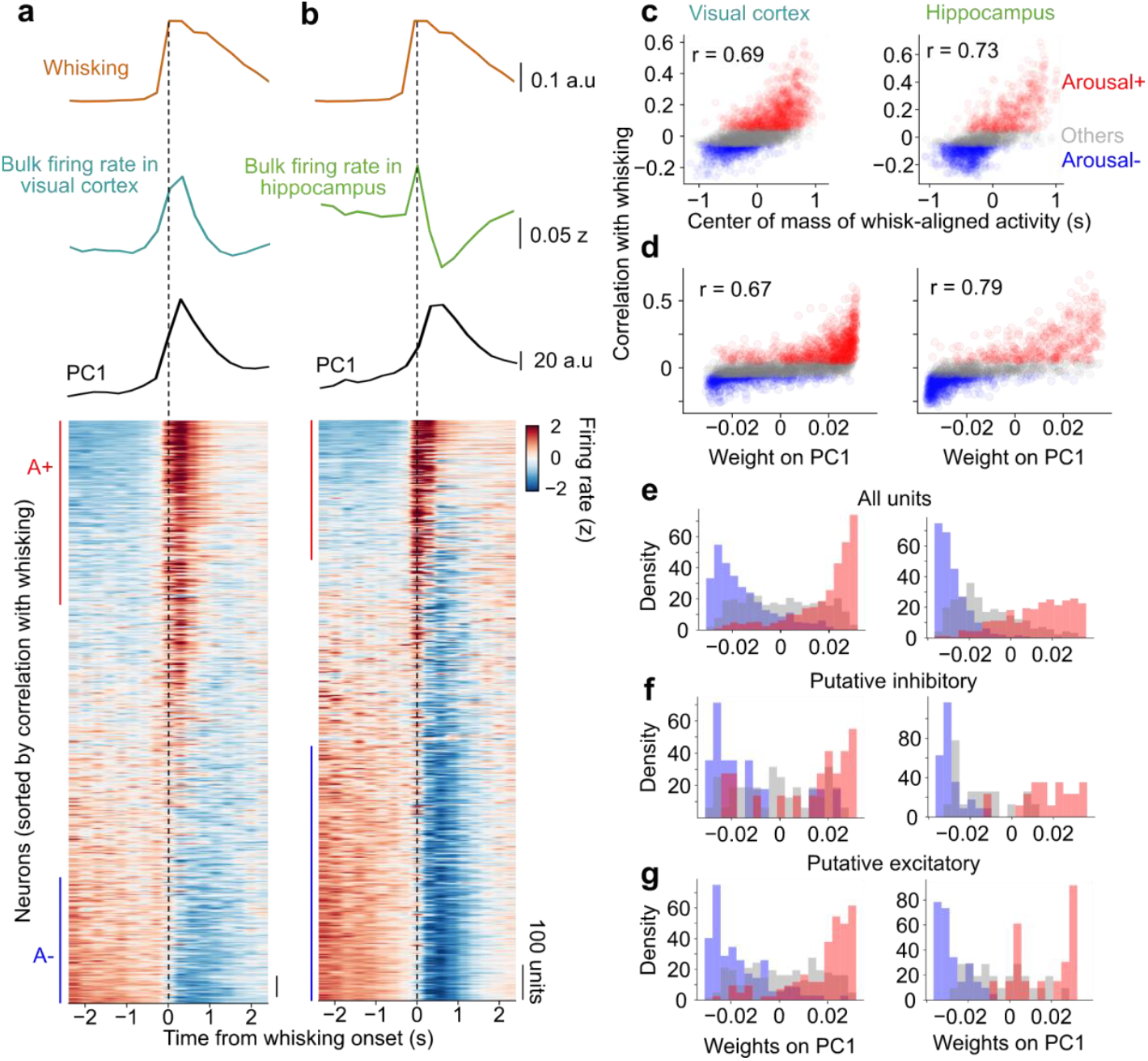
Characterization of Arousal+ and Arousal– neurons in relation to global events. **a**. This panel is reproduced from Figure 3d. Firing rate of all neurons recorded in visual cortex across all sessions, aligned to and averaged across whisking events. The traces above show the corresponding average whisking (*gold*), and average bulk firing rate (*blue*). In the matrix plot, each row/neuron is z-scored, and neurons are sorted by their correlation with whisking. We performed principal component analysis (PCA) on the z-scored whisking-evoked firing rate of all neurons. The timecourse of the first component (PC1) is shown in black. **b**. Same, for hippocampus. **c**. Correlation with whisking (used to define Arousal+ and Arousal– neurons) plotted against the centre of mass of the whisk-aligned activity (as shown in **a-b**), in visual cortex (*left*) and hippocampus (*right*). Each dot is a neuron, colour-coded by category: Arousal+ (*red*), Arousal– (*blue*) and other (*grey*) neurons. r indicates the Pearson correlation between both metrics. **d**. Correlation with whisking plotted against the weight on PC1. **e**. Histogram of neurons’ weights on PC1. Arousal+ and Arousal– neurons strongly segregated along the first component in both regions. **f**. Same as **e**, including only putative inhibitory neurons (fast-spiking single-units). **g**. Same as **e**, including only putative excitatory neurons (regular-spiking single-units). Arousal+ and Arousal– neurons are present in both putative excitatory and inhibitory populations.

**Figure S6.**
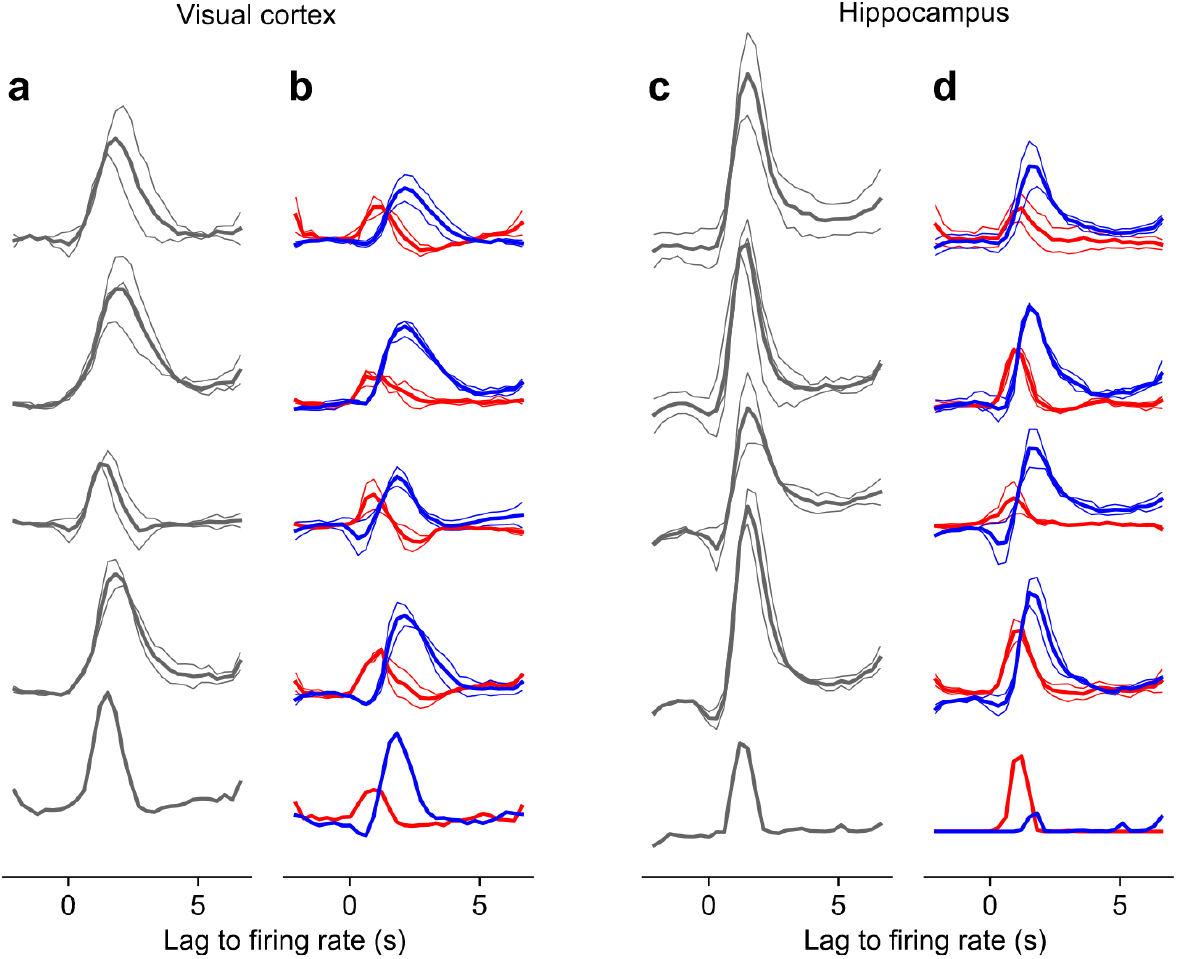
HRFs are stable across mice and sessions. **a**. Filters (HRFs) obtained to predict blood volume from bulk firing rate in visual cortex, for individual mice and recording sessions. Thin lines correspond to individual recording sessions. Thick lines correspond to average across sessions, for each mouse. Each row corresponds to a different mouse (n = 5 in total). **b**. Same with filters for Arousal+ (*red*) and Arousal– (*blue*) neurons in the combined model. **c**–**d**. Same as **a**–**b**, in hippocampus.

**Figure S7.**
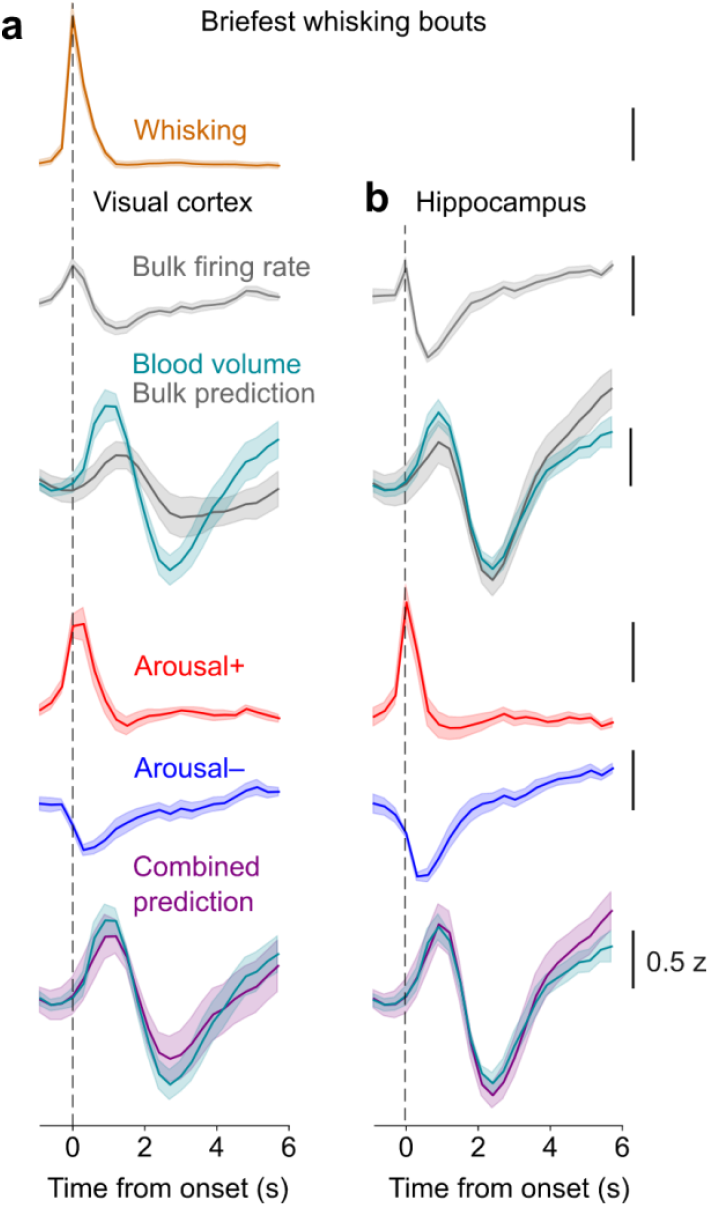
Predictions of blood volume from firing rate during briefest whisking bouts. **a**. Firing of different populations and results of blood volume predictions for very short bouts (< 1.3 s) in visual cortex. Note that since filters are fit for the entirety of each recording session, these predictions result from the same fits as in Figure 2 and Figure 3, but are assessed on different events. **b**. Same, in hippocampus.

**Figure S8.**
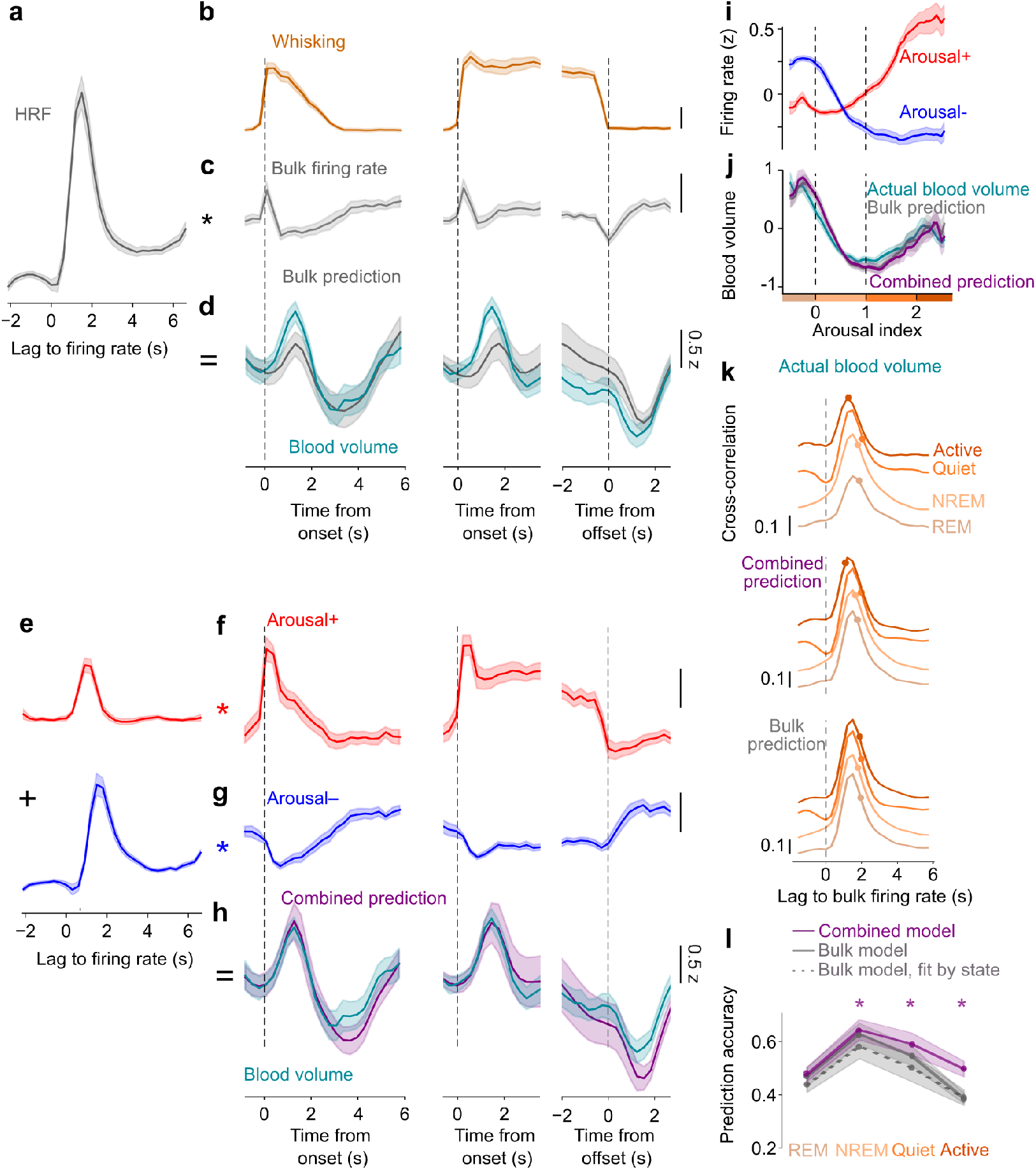
Predicting blood volume from firing rate of different populations for recordings in hippocampus. This figure replicates the analyses done in Figure 2, Figure 3 and Figure 4 for recordings in the hippocampus. **a**. The best-fitting filter to predict blood volume from average firing rate. **b**. We identified whisking bouts and looked at brief (left) and longer whisking bouts, aligned to onset (middle) or offset (right). **c**. Average changes in bulk firing rate during whisking events. **d**. Average change in blood volume during whisking events (*cyan*), and prediction from bulk firing rate (*grey*). **e**. The best-fitting filter to predict blood volume from Arousal+ and Arousal– neurons. **f**. Average changes in firing rate for Arousal+ neurons during whisking events. **g**. Same, for Arousal– neurons. **h**. Average change in blood volume during whisking events (*cyan*), and combined prediction from Arousal+ and Arousal– neurons (*purple*). **i**. Firing rate of Arousal+ (*red*) and Arousal– (*blue*) neurons as a function of arousal index, as introduced in Figure 4. **j**. Actual blood volume (*cyan*) and predictions from bulk (*grey*) and combined (*purple*) models, as a function of arousal index. **k**. Cross-correlation between bulk firing rate and actual blood volume, blood volume predicted from the bulk or combined model, for each state defined as in Figure 4. Dots indicate the centre of mass of the cross-correlations for each state. **l**. Prediction accuracy, computed as the Pearson correlation between actual and predicted blood volume depending on state. State is computed based on the arousal index in 10 s intervals. Asterisks indicate significant differences between combined and bulk prediction (paired t-test, p < 0.05, n = 10 recording sessions).

**Figure S9.**
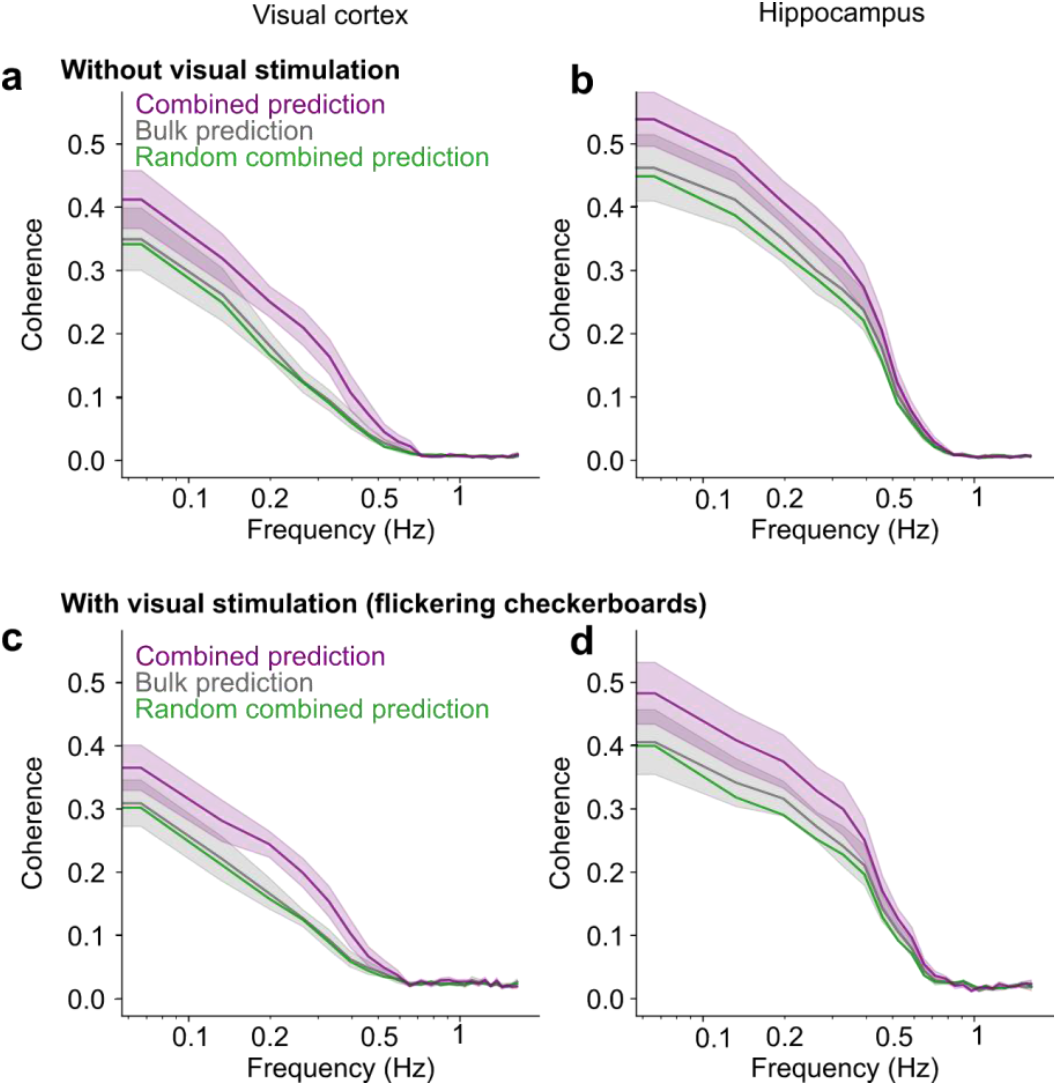
The combined model outperforms the bulk model during both spontaneous behaviour and visual stimulation. **a**. Coherence between actual and predicted blood volume in visual cortex, using only spontaneous activity (grey screen). The combined model (*purple*) outperforms the bulk model (*grey*) (p = 2e-3). **b**. Same in hippocampus (p = 3e-4). **c**–**d**, Same as a–b, using only visual stimulation epochs (flickering checkerboards). The combined model also outperforms the bulk in this condition in both regions (p = 0.014 in visual cortex, p = 6e-3 in hippocampus).

**Figure S10.**
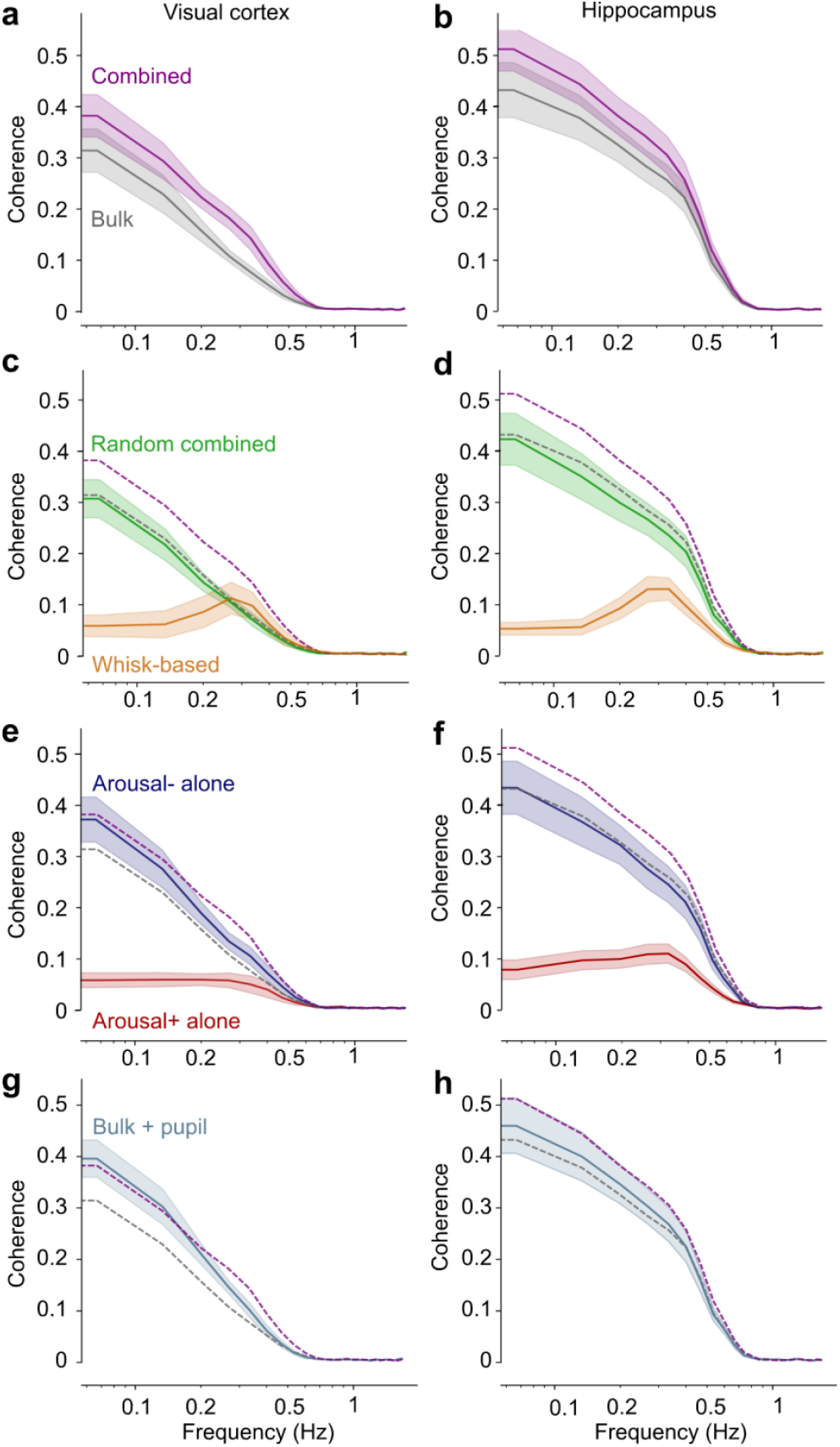
Prediction of blood volume with alternative models. **a**. For reference, coherence between actual and predicted blood volume for the predictions from bulk firing rate (*grey*) or from Arousal+ and Arousal– neurons (*purple*), in visual cortex (reproduced from Figure 3j). Shaded area shows the s.e. across sessions. **b**. Same, for recordings in hippocampus. **c**. To test whether the improved performance of the combined model stemmed from more degrees of freedom in the prediction, we computed predictions of blood volume in visual cortex by fitting filters for the average firing rate of two random populations of neurons, matched in size with Arousal+ and Arousal– populations (random combined prediction, *green*). This prediction led to a coherence with actual blood volume similar to the bulk prediction (*dashed grey*, p = 0.20 across frequencies), and worse than the combined prediction (*dashed purple*, p = 2e-4). Predictions from whisking alone (*gold*) were also worse than the combined prediction (p = 4e-5). **d**. The same results were found in hippocampus: the predictions from the two random populations were worse than the bulk (p = 0.02) and combined (p = 2e-5) predictions, as for the prediction from whisking alone (p = 4e-5). **e**. To test whether the improved performance of the combined model stemmed from more homogeneous neural populations, we computed predictions from either Arousal+ or Arousal– firing rate alone. Both led to significantly worse predictions than the combined model (p = 1e-5 for Arousal+, p = 9e-3 for Arousal–). **f**. The same result was found in hippocampus (p = 9e-5 for Arousal+, p= 3e-4 for Arousal–). All p-values result from paired t-tests between the average coherence for frequencies < 0.7 Hz, across sessions (n = 10 independent recording sessions per region). **g**. We also fit a dual-HRF model based on bulk firing rate and pupil size. The resulting predictions were better than bulk predictions (p = 7e-3) and similar to our combined model with Arousal+ and Arousal-neurons (p = 0.35). **h**. In hippocampus, the same model did not show improvement over the bulk model (p = 0.09) and performed worse than the combined model (p = 3e-3).

**Figure S11.**
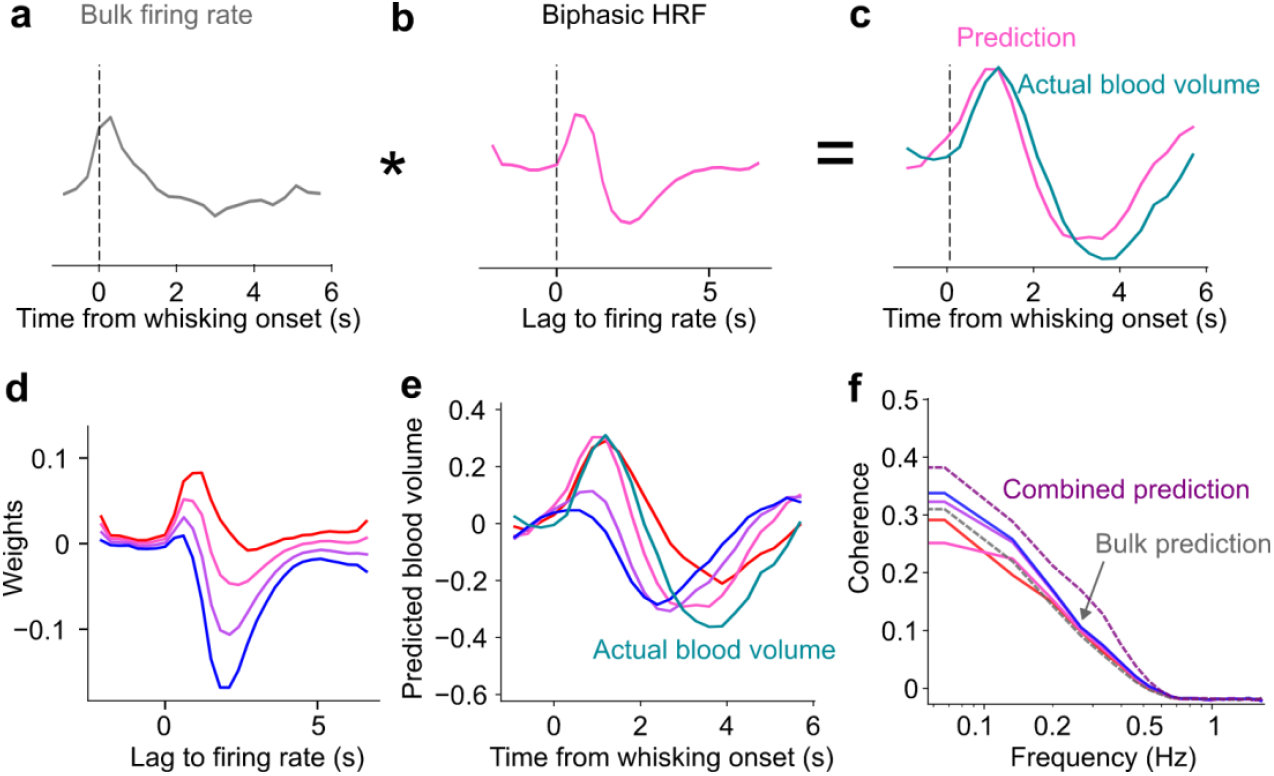
Prediction of blood volume from average firing rate with biphasic filters. We tested whether a biphasic filter on bulk firing rate could produce similar results as those obtained with the combined prediction. We convolved bulk firing rate (**a**) with a biphasic filter (**b**) obtained by taking a weighted difference between the filter obtained for Arousal+ neurons and the filter for Arousal– neurons in the combined prediction. We then aligned the obtained predictions to whisking onsets and compared them to actual blood volume (**c**). **d**. We used this procedure with different relative weights between Arousal+ and Arousal– filters. **e**. None of these filters recapitulated the dynamics seen in blood volume (*cyan*) following whisking events. Some produced biphasic responses but with a different timescale and amplitude. **f**. Coherence between actual and predicted blood volume for the different filters. All the predictions obtained through this procedure were significantly worse than the combined prediction (*dashed purple*, p < 0.02 in visual cortex, shown here, p < 2e-3 in hippocampus), and similar or worse than the bulk prediction (*dashed grey*, p = 0.002–0.78 in visual cortex, p = 0.02–0.22 in hippocampus).

**Figure S12.**
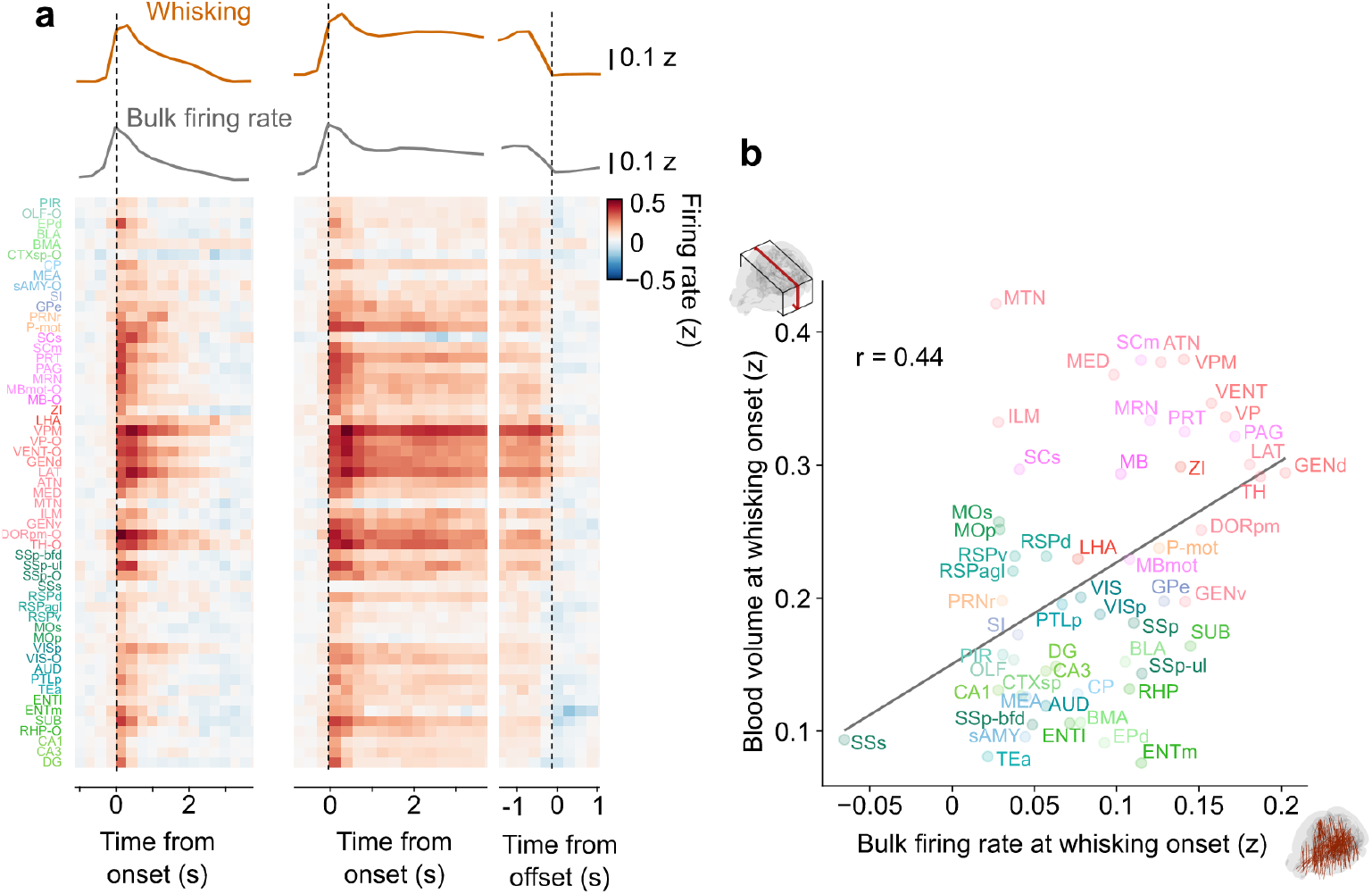
Brainwide bulk firing rate during whisking events. **a**. Average firing rate during brief whisking events (*left*), longer whisking events, aligned to onset (*middle*) or offset *(right*), for all regions. The grey trace indicates the average across all regions. **b**. Average blood volume (from the brainwide fUSI recordings) 1.2–1.8 s after whisking onset plotted against average bulk firing rate (from the brainwide Neuropixels recordings) 0–0.6 s after whisking onset. Each dot is a brain region, accompanied by its Allen acronym. The line shows a linear fit and r indicates the Pearson correlation.

**Figure S13.**
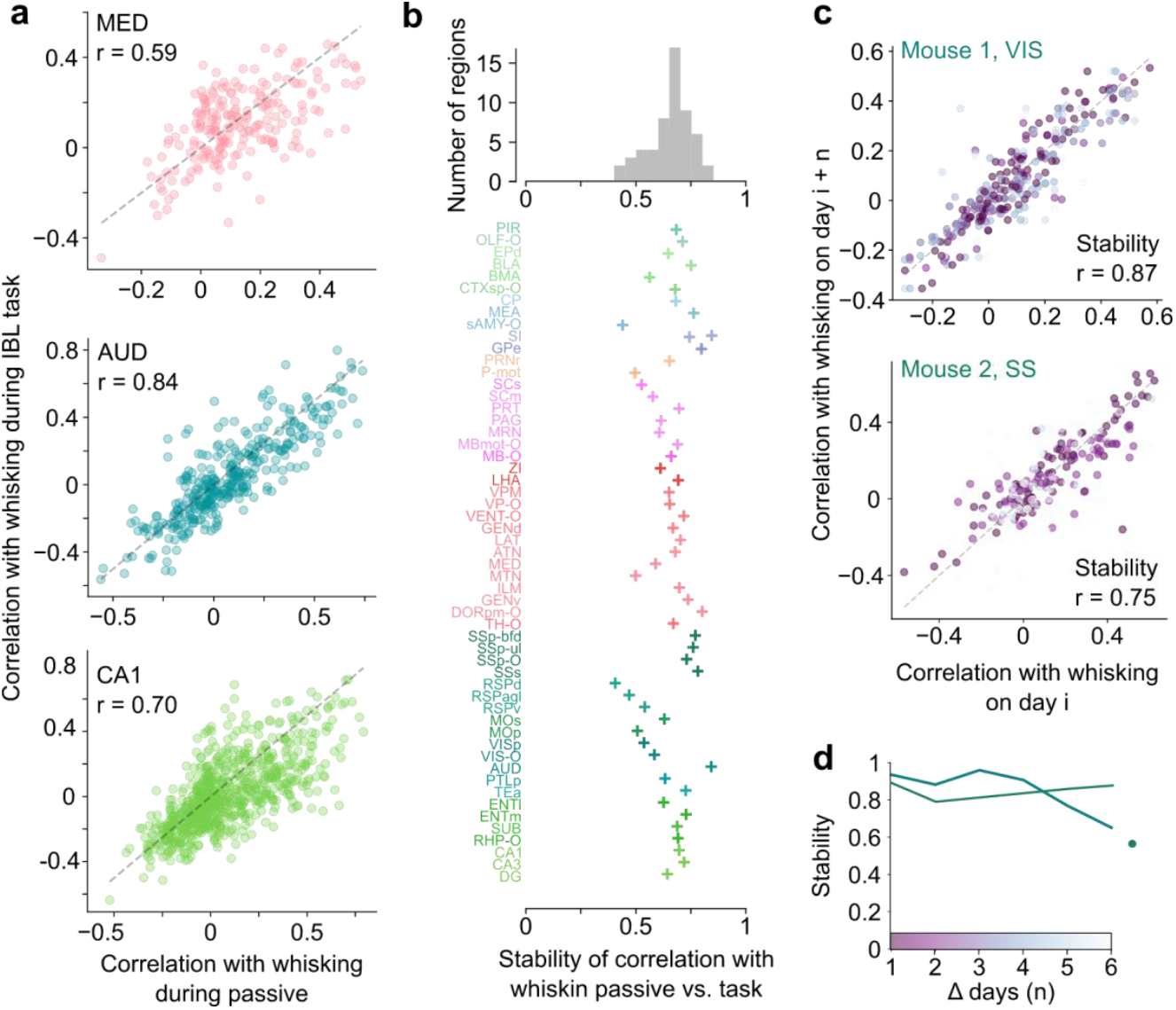
Stability of correlation with whisking across behavioural contexts and days. **a**. To assess the stability of our populations across behavioural contexts, we computed the correlation between neurons’ firing rate and whisking in the IBL dataset while mice were engaged in the task. For three example regions, we plotted this correlation against the correlation during passive epochs, which we used to define Arousal+ and Arousal– populations. Each dot is a neuron. r indicates the Pearson correlation, which we thereafter refer to as ‘stability’. **b**. *Top*: histogram of stability per brain region, defined as the correlation between the correlation with whisking in passive conditions vs. in the task (as shown in **a**). *Bottom*: Stability values for all brain regions. **c**. To assess the stability of our populations across days, we chronically recorded neurons in two mice (mouse 1 in visual cortex, mouse 2 in somatosensory cortex) and tracked them across days. Here, we compare the correlation with whisking for matched neurons recorded on different sessions separated from 1 to 6+ days (colour-coded). Each dot is a pair of matched neurons across days. We define stability as the correlation between these points. **d**. Average stability for different time intervals between recording sessions. The rightmost point indicates the stability for pairs with intervals longer than 6 days (ranging from 7 to 19 days), for mouse 2.

**Figure S14.**
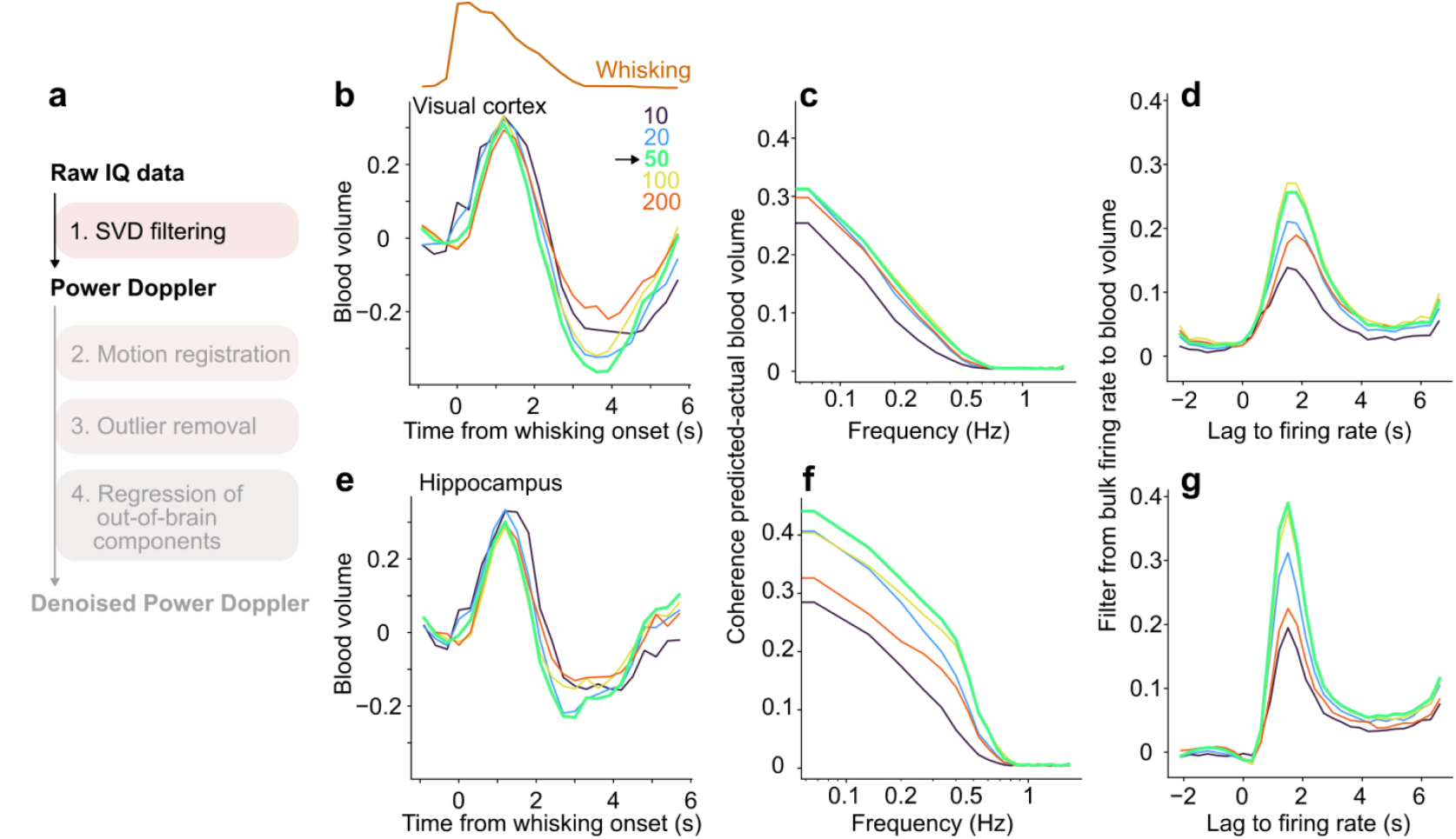
Varying number of components removed in the SVD. **a**. Summary of pre-processing and denoising pipeline. Here we focus on step 1, which relies on filtering the raw complex IQ data through singular value decomposition (SVD) on batches of 300 images (600 ms at 500 Hz). The first n components, representing coherent tissue motion, are subtracted from the signal and the squared modulus of the signal is taken to obtain Power Doppler. Here we varied the number of components removed (n) and examined the effects on our main results. **b**. Average whisking-related change in blood volume in visual cortex depending on n. **c**. Coherence between blood volume measured and predicted from bulk firing rate, depending on n. A different model is fit for each value of n. **d**. Best-fitting filter to predict blood volume from bulk firing rate depending on n. **e**–**g**. Same as **b–d**, in hippocampus. In both regions, the best prediction from neural activity is obtained for n = 50, and stable between 20–100. Removing less components leads to the appearance of artefacts synchronous with motion (see t = 0 s in panel **a, e**). Removing more components leads to a loss in predictability from neural activity, presumably because true signal is removed. In the rest of the paper, we always use n = 50.

**Figure S15.**
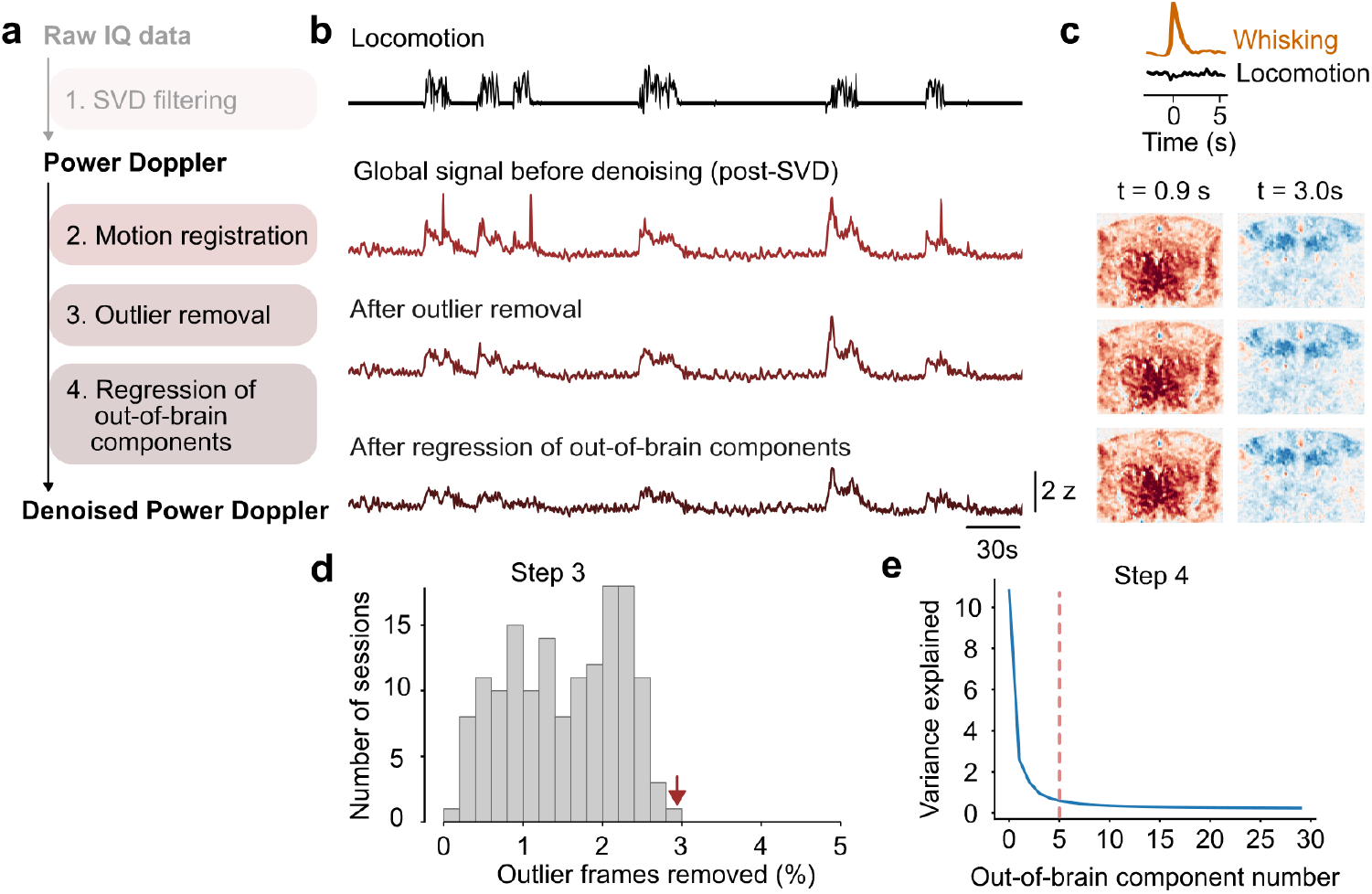
Effects of denoising steps. **a**. Summary of pre-processing and denoising pipeline. Here we focus on steps 3-4. **b**. Example snippet from a recording session showing the global signal, i.e. the average of all z-scored brain voxels after each denoising step, along with locomotion. The steps particularly affect epochs of locomotion, which tend to show more motion artefacts. **c**. For the example recording session in b, we extracted whisking events in the absence of locomotion (as in Figure 1). Bottom plots show the average signal for all voxels, 0.9 s and 3.0 s after whisking onset (recentred by the pre-whisking baseline), after each denoising step. The different steps have little effect on the whisking-related changes in the absence of locomotion, which are weakly affected by motion artefacts. **d**. Distribution of the number of outlier frames removed in step 3 across all recording sessions. The arrow indicates the position of the example session shown in b-c. **e**. Variance explained by each out-of-brain component, as used for regression in step 4. The dotted line indicates the number of components we chose to keep as regressors.

## References

1 Heeger, D. J. & Ress, D. What does fMRI tell us about neuronal activity? Nat Rev Neurosci 3, 142–151 (2002). 10.1038/nrn730

2 Drew, P. J. Vascular and neural basis of the BOLD signal. Curr Opin Neurobiol 58, 61–69 (2019). 10.1016/j.conb.2019.06.004

3 Logothetis, N. K. & Wandell, B. A. Interpreting the BOLD signal. Annu Rev Physiol 66, 735–769 (2004). 10.1146/annurev.physiol.66.082602.092845

4 Hillman, E. M. Coupling mechanism and significance of the BOLD signal: a status report. Annu Rev Neurosci 37, 161–181 (2014). 10.1146/annurev-neuro-071013-014111

5 Drew, P. J. Neurovascular coupling: motive unknown. Trends Neurosci 45, 809–819 (2022). 10.1016/j.tins.2022.08.004

6 Logothetis, N. K., Pauls, J., Augath, M., Trinath, T. & Oeltermann, A. Neurophysiological investigation of the basis of the fMRI signal. Nature 412, 150–157 (2001). 10.1038/35084005

7 Devor, A. et al. Coupling of the cortical hemodynamic response to cortical and thalamic neuronal activity. PNAS 102, 3822–3827 (2005). 10.1073/pnas.0407789102

8 Mukamel, R. et al. Coupling Between Neuronal Firing, Field Potentials, and fMRI in Human Auditory Cortex. Science 309, 951–954 (2005). 10.1126/science.1110913

9 Shmuel, A., Augath, M., Oeltermann, A. & Logothetis, N. K. Negative functional MRI response correlates with decreases in neuronal activity in monkey visual area V1. Nature Neuroscience 9, 569–577 (2006). 10.1038/nn1675

10 Nunez-Elizalde, A. O. et al. Neural correlates of blood flow measured by ultrasound. Neuron 110, 1631–1640 e1634 (2022). 10.1016/j.neuron.2022.02.012

11 Sirotin, Y. B. & Das, A. Anticipatory haemodynamic signals in sensory cortex not predicted by local neuronal activity. Nature 457, 475–479 (2009). 10.1038/nature07664

12 Turner, K. L., Gheres, K. W. & Drew, P. J. Relating Pupil Diameter and Blinking to Cortical Activity and Hemodynamics across Arousal States. J Neurosci 43, 949–964 (2023). 10.1523/JNEUROSCI.1244-22.2022

13 Rauscher, C. B. et al. Neurovascular Impulse Response Function (IRF) during spontaneous activity differentially reflects intrinsic neuromodulation across cortical regions. BioRxiv (2024). 10.1101/2024.09.14.612514

14 Ekstrom, A. How and when the fMRI BOLD signal relates to underlying neural activity: the danger in dissociation. Brain Res Rev 62, 233–244 (2010). 10.1016/j.brainresrev.2009.12.004

15 Jukovskaya, N., Tiret, P., Lecoq, J. & Charpak, S. What does local functional hyperemia tell about local neuronal activation? J Neurosci 31, 1579–1582 (2011). 10.1523/JNEUROSCI.3146-10.2011

16 Devonshire, I. M. et al. Neurovascular coupling is brain region-dependent. Neuroimage 59, 1997–2006 (2012). 10.1016/j.neuroimage.2011.09.050

17 Huo, B. X., Smith, J. B. & Drew, P. J. Neurovascular coupling and decoupling in the cortex during voluntary locomotion. J Neurosci 34, 10975–10981 (2014). 10.1523/JNEUROSCI.1369-14.2014

18 Devor, A. et al. Stimulus-induced changes in blood flow and 2-deoxyglucose uptake dissociate in ipsilateral somatosensory cortex. J Neurosci 28, 14347–14357 (2008). 10.1523/JNEUROSCI.4307-08.2008

19 Shih, Y. Y. et al. A new scenario for negative functional magnetic resonance imaging signals: endogenous neurotransmission. J Neurosci 29, 3036–3044 (2009). 10.1523/JNEUROSCI.3447-08.2009

20 Boorman, L. et al. Negative blood oxygen level dependence in the rat: a model for investigating the role of suppression in neurovascular coupling. J Neurosci 30, 4285–4294 (2010). 10.1523/JNEUROSCI.6063-09.2010

21 Scholvinck, M. L., Maier, A., Ye, F. Q., Duyn, J. H. & Leopold, D. A. Neural basis of global resting-state fMRI activity. PNAS 107, 10238–10243 (2010). 10.1073/pnas.0913110107

22 Meyer-Baese, L. et al. Cortical Networks Relating to Arousal Are Differentially Coupled to Neural Activity and Hemodynamics. J Neurosci 44 (2024). 10.1523/JNEUROSCI.0298-23.2024

23 Winder, A. T., Echagarruga, C., Zhang, Q. & Drew, P. Weak correlations between hemodynamic signals and ongoing neural activity during the resting state. Nat Neurosci 20, 1761–1769 (2017). 10.1038/s41593-017-0007-y

24 Turner, K. L., Gheres, K. W., Proctor, E. A. & Drew, P. Neurovascular coupling and bilateral connectivity during NREM and REM sleep. eLife 9 (2020). 10.7554/eLife.62071

25 Boynton, G. M. Spikes, BOLD, attention, and awareness: a comparison of electrophysiological and fMRI signals in V1. J Vis 11, 12 (2011). 10.1167/11.5.12

26 Cardoso, M. M., Sirotin, Y. B., Lima, B., Glushenkova, E. & Das, A. The neuroimaging signal is a linear sum of neurally distinct stimulus- and task-related components. Nat Neurosci 15, 1298–1306 (2012). 10.1038/nn.3170

27 Nir, Y. et al. Coupling between neuronal firing rate, gamma LFP, and BOLD fMRI is related to interneuronal correlations. Curr Biol 17, 1275–1285 (2007). 10.1016/j.cub.2007.06.066

28 Buzsaki, G., Kaila, K. & Raichle, M. Inhibition and brain work. Neuron 56, 771–783 (2007). 10.1016/j.neuron.2007.11.008

29 Howarth, C., Mishra, A. & Hall, C. N. More than just summed neuronal activity: how multiple cell types shape the BOLD response. Philos Trans R Soc Lond B Biol Sci 376, 20190630 (2021). 10.1098/rstb.2019.0630

30 Attwell, D. et al. Glial and neuronal control of brain blood flow. Nature 468, 232–243 (2010). 10.1038/nature09613

31 Lee, L. et al. Key Aspects of Neurovascular Control Mediated by Specific Populations of Inhibitory Cortical Interneurons. Cerebral Cortex 30, 2452–2464 (2020). 10.1093/cercor/bhz251

32 Ruff, C. F. et al. Long-range inhibitory neurons mediate cortical neurovascular coupling. Cell Rep 43, 113970 (2024). 10.1016/j.celrep.2024.113970

33 Turner, K. L. et al. Type-I nNOS neurons orchestrate cortical neural activity and vasomotion. bioRxiv (2025). 10.1101/2025.01.21.634042

34 Hauglund, L. N. et al. Norepinephrine-mediated slow vasomotion drives glymphatic clearance during sleep. Cell 188, 606–622.e617 (2025). 10.1016/j.cell.2024.11.027

35 Zaldivar, D. et al. Brain-wide functional connectivity of face patch neurons during rest. PNAS 119, e2206559119 (2022). 10.1073/pnas.2206559119

36 Macé, E. et al. Functional ultrasound imaging of the brain. Nat Methods 8, 662–664 (2011). 10.1038/nmeth.1641

37 Garcia-Junco-Clemente, P. et al. An inhibitory pull– push circuit in frontal cortex. Nat Neurosci 20, 389–392 (2017). 10.1038/nn.4483

38 Stringer, C. et al. Spontaneous behaviors drive multidimensional, brainwide activity. Science 364, 255 (2019). 10.1126/science.aav7893

39 Brécier, A., Borel, M., Urbain, N. & Gentet, J. L. Vigilance and Behavioral State-Dependent Modulation of Cortical Neuronal Activity throughout the Sleep/Wake Cycle. J Neurosci 42, 4852–4866 (2022). 10.1523/JNEUROSCI.1400-21.2022

40 International Brain Laboratory et al. A brain-wide map of neural activity during complex behaviour. Nature 645, 177–191 (2025). 10.1038/s41586-025-09235-0

41 Jun, J. J. et al. Fully integrated silicon probes for high-density recording of neural activity. Nature 551, 232–236 (2017). 10.1038/nature24636

42 Wang, Q. et al. The Allen Mouse Brain Common Coordinate Framework: A 3D Reference Atlas. Cell 181, 936–953.e920 (2020). 10.1016/j.cell.2020.04.007

43 Drew, P. J., Winder, A. T. & Zhang, Q. Twitches, Blinks, and Fidgets: Important Generators of Ongoing Neural Activity. Neuroscientist 25, 298–313 (2019). 10.1177/1073858418805427

44 Collins, L., Francis, J., Emanuel, B. & McCormick, D. A. Cholinergic and noradrenergic axonal activity contains a behavioral-state signal that is coordinated across the dorsal cortex. eLife 12, e81826 (2023). 10.7554/eLife.81826

45 McGinley, M. J., David, S. V. & McCormick, D. A. Cortical Membrane Potential Signature of Optimal States for Sensory Signal Detection. Neuron 87, 179–192 (2015). 10.1016/j.neuron.2015.05.038

46 Ma, Y. et al. Resting-state hemodynamics are spatiotemporally coupled to synchronized and symmetric neural activity in excitatory neurons. PNAS 113, E8463–E8471 (2016). 10.1073/pnas.1525369113

47 Pisauro, M. A., Benucci, A. & Carandini, M. Local and global contributions to hemodynamic activity in mouse cortex. J Neurophysiol 115, 2931–2936 (2016). 10.1152/jn.00125.2016

48 Aydin, A.-K. et al. Transfer functions linking neural calcium to single voxel functional ultrasound signal. Nat Comm 11, 2954 (2020). 10.1038/s41467-020-16774-9

49 Liu, X., Leopold, D. A. & Yang, Y. Single-neuron firing cascades underlie global spontaneous brain events. PNAS 118 (2021). 10.1073/pnas.2105395118

50 Yang, Y., Leopold, D. A., Duyn, J. H., Sipe, G. O. & Liu, X. Sensory Encoding Alternates With Hippocampal Ripples across Cycles of Forebrain Spiking Cascades. Adv Sci (Weinh) 12, e2406224 (2025). 10.1002/advs.202406224

51 Reimer, J. et al. Pupil Fluctuations Track Fast Switching of Cortical States during Quiet Wakefulness. Neuron 84, 355–362 (2014). 10.1016/j.neuron.2014.09.033

52 Vinck, M., Batista-Brito, R., Knoblich, U. & Cardin, J. A. Arousal and locomotion make distinct contributions to cortical activity patterns and visual encoding. Neuron 86, 740–754 (2015). 10.1016/j.neuron.2015.03.028

53 Reimer, J. et al. Pupil fluctuations track rapid changes in adrenergic and cholinergic activity in cortex. Nat Comm 7, 13289 (2016). 10.1038/ncomms13289

54 Pais-Roldán, P. et al. Indexing brain state-dependent pupil dynamics with simultaneous fMRI and optical fiber calcium recording. PNAS 117, 6875–6882 (2020). 10.1073/pnas.1909937117

55 McGinley, M. J. et al. Waking State: Rapid Variations Modulate Neural and Behavioral Responses. Neuron 87, 1143–1161 (2015). 10.1016/j.neuron.2015.09.012

56 Yuzgec, O., Prsa, M., Zimmermann, R. & Huber, D. Pupil Size Coupling to Cortical States Protects the Stability of Deep Sleep via Parasympathetic Modulation. Curr Biol 28, 392–400 e393 (2018). 10.1016/j.cub.2017.12.049

57 Fulda, S. et al. Rapid eye movements during sleep in mice: High trait-like stability qualifies rapid eye movement density for characterization of phenotypic variation in sleep patterns of rodents. BMC Neuroscience 12, 110 (2011). 10.1186/1471-2202-12-110

58 Sanchez-Lopez, A. & Escudero, M. Tonic and phasic components of eye movements during REM sleep in the rat. Eur J Neurosci 33, 2129–2138 (2011). 10.1111/j.1460-9568.2011.07702.x

59 Bergel, A., Deffieux, T., Demene, C., Tanter, M. & Cohen, I. Local hippocampal fast gamma rhythms precede brain-wide hyperemic patterns during spontaneous rodent REM sleep. Nat Comm 9, 5364 (2018). 10.1038/s41467-018-07752-3

60 Bimbard, C. et al. An adaptable, reusable, and light implant for chronic Neuropixels probes. eLife 13 (2025). 10.7554/eLife.98522

61 van Beest, E. H. et al. Tracking neurons across days with high-density probes. Nat Methods 22, 778–787 (2025). 10.1038/s41592-024-02440-1

62 Steinmetz, N. A., Zatka-Haas, P., Carandini, M. & Harris, K. D. Distributed coding of choice, action and engagement across the mouse brain. Nature 576, 266–273 (2019). 10.1038/s41586-019-1787-x

63 Allen, W. E. et al. Thirst regulates motivated behavior through modulation of brainwide neural population dynamics. Science 364, 253 (2019). 10.1126/science.aav3932

64 Shimaoka, D., Harris, D. K. & Carandini, M. Effects of Arousal on Mouse Sensory Cortex Depend on Modality. Cell Rep 22, 3160–3167 (2018). 10.1016/j.celrep.2018.02.092

65 Musall, S., Kaufman, M. T., Juavinett, A. L., Gluf, S. & Churchland, A. K. Single-trial neural dynamics are dominated by richly varied movements. Nat Neurosci 22, 1677–1686 (2019). 10.1038/s41593-019-0502-4

66 Han, Z. et al. Awake and behaving mouse fMRI during Go/No-Go task. Neuroimage 188, 733–742 (2019). 10.1016/j.neuroimage.2019.01.002

67 Zhang, J. & Northoff, G. Beyond noise to function: reframing the global brain activity and its dynamic topography. Commun Biol 5, 1350 (2022). 10.1038/s42003-022-04297-6

68 Ma, Y., Ma, Z., Liang, Z., Neuberger, T. & Zhang, N. Global brain signal in awake rats. Brain Struct Funct 225, 227–240 (2020). 10.1007/s00429-019-01996-5

69 Gutierrez-Barragan, D. et al. Unique spatiotemporal fMRI dynamics in the awake mouse brain. Curr Biol 32, 631–644 e636 (2022). 10.1016/j.cub.2021.12.015

70 Hamada, H. T. et al. Optogenetic activation of dorsal raphe serotonin neurons induces brain-wide activation. Nat Comm 15, 4152 (2024). 10.1038/s41467-024-48489-6

71 Chang, C. et al. Tracking brain arousal fluctuations with fMRI. PNAS 113, 4518–4523 (2016). 10.1073/pnas.1520613113

72 Turchi, J. et al. The Basal Forebrain Regulates Global Resting-State fMRI Fluctuations. Neuron 97, 940–952 e944 (2018). 10.1016/j.neuron.2018.01.032

73 Wong, C. W., Olafsson, V., Tal, O. & Liu, T. T. The amplitude of the resting-state fMRI global signal is related to EEG vigilance measures. Neuroimage 83, 983–990 (2013). 10.1016/j.neuroimage.2013.07.057

74 Yellin, D., Berkovich-Ohana, A. & Malach, R. Coupling between pupil fluctuations and resting-state fMRI uncovers a slow build-up of antagonistic responses in the human cortex. Neuroimage 106, 414–427 (2015). 10.1016/j.neuroimage.2014.11.034

75 Breeden, A. L., Siegle, G. J., Norr, M. E., Gordon, E. M. & Vaidya, C. J. Coupling between spontaneous pupillary fluctuations and brain activity relates to inattentiveness. Eur J Neurosci 45, 260–266 (2017). 10.1111/ejn.13424

76 Schneider, M. et al. Spontaneous pupil dilations during the resting state are associated with activation of the salience network. Neuroimage 139, 189–201 (2016). 10.1016/j.neuroimage.2016.06.011

77 Raut, R. V. et al. Global waves synchronize the brain’s functional systems with fluctuating arousal. Science Advances 7, eabf2709 (2021). 10.1126/sciadv.abf2709

78 Bolt, T. et al. Autonomic physiological coupling of the global fMRI signal. Nat Neurosci 28, 1327–1335 (2025). 10.1038/s41593-025-01945-y

79 Roth, Z. N., Ryoo, M. & Merriam, E. P. Task-related activity in human visual cortex. PLoS Biol 18, e3000921 (2020). 10.1371/journal.pbio.3000921

80 Setzer, B. et al. A temporal sequence of thalamic activity unfolds at transitions in behavioral arousal state. Nat Comm 13, 5442 (2022). 10.1038/s41467-022-33010-8

81 Fultz, N. E. et al. Coupled electrophysiological, hemodynamic, and cerebrospinal fluid oscillations in human sleep. Science 366, 628–631 (2019). 10.1126/science.aax5440

82 Fisher, P. S. et al. Stereotypic wheel running decreases cortical activity in mice. Nat Comm 7, 13138 (2016). 10.1038/ncomms13138

83 Christensen, J. A. & Pillow, W. J. Reduced neural activity but improved coding in rodent higher-order visual cortex during locomotion. Nat Comm 13 (2022). 10.1038/s41467-022-29200-z

84 Schneider, M. D., Nelson, A. & Mooney, R. A synaptic and circuit basis for corollary discharge in the auditory cortex. Nature 513, 189–194 (2014). 10.1038/nature13724

85 Bergel, A. et al. Adaptive modulation of brain hemodynamics across stereotyped running episodes. Nat Comm 11, 6193 (2020). 10.1038/s41467-020-19948-7

86 Ratliff, J. M. et al. Neocortical long-range inhibition promotes cortical synchrony and sleep. bioRxiv (2024). 10.1101/2024.06.20.599756

87 Perrenoud, Q. et al. Characterization of Type I and Type II nNOS-Expressing Interneurons in the Barrel Cortex of Mouse. Front Neural Circuits 6, 36 (2012). 10.3389/fncir.2012.00036

88 Echagarruga, T. C., Gheres, W. K., Norwood, N. J. & Drew, J. P. nNOS-expressing interneurons control basal and behaviorally evoked arterial dilation in somatosensory cortex of mice. eLife 9 (2020). 10.7554/eLife.60533

89 Valero, M. et al. Sleep down state-active ID2/Nkx2.1 interneurons in the neocortex. Nat Neurosci 24, 401–411 (2021). 10.1038/s41593-021-00797-6

90 Bugeon, S. et al. A transcriptomic axis predicts state modulation of cortical interneurons. Nature 607, 330–338 (2022). 10.1038/s41586-022-04915-7

91 McCormick, D.A. Cholinergic and noradrenergic modulation of thalamocortical processing. Trends in neurosciences 12, 215–221 (1989). 10.1016/0166-2236(89)90125-2

92 Eggermann, E., Kremer, Y., Crochet, S. & Petersen, C. C. H. Cholinergic signals in mouse barrel cortex during active whisker sensing. Cell Rep 9, 1654–1660 (2014). 10.1016/j.celrep.2014.11.005

93 Chen, N., Sugihara, H. & Sur, M. An acetylcholine-activated microcircuit drives temporal dynamics of cortical activity. Nat Neurosci 18, 892–902 (2015). 10.1038/nn.4002

94 Inácio, R. A. et al. Brain-wide presynaptic networks of functionally distinct cortical neurons. Nature (2025). 10.1038/s41586-025-08631-w

95 Cauli, B. et al. Cortical GABA Interneurons in Neurovascular Coupling: Relays for Subcortical Vasoactive Pathways. J Neurosci 24, 8940–8949 (2004). 10.1523/JNEUROSCI.3065-04.2004

96 Gentet, J. L., Avermann, M., Matyas, F., Staiger, F. J. & Petersen, C. H. C. Membrane Potential Dynamics of GABAergic Neurons in the Barrel Cortex of Behaving Mice. Neuron 65, 422–435 (2010). 10.1016/j.neuron.2010.01.006

97 Gentet, L. J. et al. Unique functional properties of somatostatin-expressing GABAergic neurons in mouse barrel cortex. Nat Neurosci 15, 607–612 (2012). 10.1038/nn.3051

98 Fu, Y. et al. A cortical circuit for gain control by behavioral state. Cell 156, 1139–1152 (2014). 10.1016/j.cell.2014.01.050

99 Muñoz, W., Tremblay, R., Levenstein, D. & Rudy, B. Layer-specific modulation of neocortical dendritic inhibition during active wakefulness. Science 355, 954–959 (2017). 10.1126/science.aag2599

100 Mayerich, D., Abbott, L. & McCormick, B. Knife-edge scanning microscopy for imaging and reconstruction of three-dimensional anatomical structures of the mouse brain. Journal of Microscopy 231, 134–143 (2008). 10.1111/j.1365-2818.2008.02024.x

101 Ragan, T. et al. Serial two-photon tomography for automated ex vivo mouse brain imaging. Nat Methods 9, 255–258 (2012). 10.1038/nmeth.1854

102 Demené, C. et al. Spatiotemporal Clutter Filtering of Ultrafast Ultrasound Data Highly Increases Doppler and fUltrasound Sensitivity. IEEE Transactions on Medical Imaging 34, 2271–2285 (2015). 10.1109/TMI.2015.2428634

103 El Hady, A. et al. Chronic brain functional ultrasound imaging in freely moving rodents performing cognitive tasks. J Neurosci Methods 403, 110033 (2024). 10.1016/j.jneumeth.2023.110033

104 Tiran, E. et al. Transcranial Functional Ultrasound Imaging in Freely Moving Awake Mice and Anesthetized Young Rats without Contrast Agent. Ultrasound Med Biol 43, 1679–1689 (2017). 10.1016/j.ultrasmedbio.2017.03.011

105 Pnevmatikakis, A. E. & Giovannucci, A. NoRMCorre: An online algorithm for piecewise rigid motion correction of calcium imaging data. J Neurosci Methods 291, 83–94 (2017). 10.1016/j.jneumeth.2017.07.031

106 Tyson, L. A. et al. Accurate determination of marker location within whole-brain microscopy images. Sci Rep 12 (2022). 10.1038/s41598-021-04676-9

107 Niedworok, J. C. et al. aMAP is a validated pipeline for registration and segmentation of high-resolution mouse brain data. Nat Comm 7, 11879 (2016). 10.1038/ncomms11879

108 Claudi, F. et al. BrainGlobe Atlas API: a common interface for neuroanatomical atlases. Journal of Open Source Software 5, 2668 (2020). 10.21105/joss.02668

109 Mathis, A. et al. DeepLabCut: markerless pose estimation of user-defined body parts with deep learning. Nat Neurosci 21, 1281–1289 (2018). 10.1038/s41593-018-0209-y

